# Preclinical Efficacy of Tasquinimod in Myelodysplastic Neoplasms: Restoring Erythropoiesis and Mitigating Bone Loss

**DOI:** 10.1101/2025.07.01.660147

**Authors:** Manja Wobus, Heike Weidner, Rebekka Wehner, Anna-Lena Baumann, Kristin Möbus, Ekaterina Balaian, Marie Törngren, Erik Vahtola, Helena Eriksson, Susann Winter, Uwe Platzbecker, Triantafyllos Chavakis, Lorenz Christian Hofbauer, Martina Rauner, Martin Bornhäuser, Katja Sockel

**Author notes:** These authors contributed equally to this work (Manja Wobus, Heike Weidner).

## Abstract

Myelodysplastic neoplasms (MDS) are clonal disorders characterized by ineffective hematopoiesis, dysplasia, and a risk of transformation into acute myeloid leukemia. MDS is also associated with a higher incidence of osteoporosis, suggesting a complex interplay between hematopoiesis, the bone marrow (BM) microenvironment, and bone homeostasis. Targeting inflammation has emerged as a promising therapeutic strategy, particularly in lower-risk MDS. Tasquinimod (TASQ) is a small-molecule inhibitor of the inflammatory alarmin S100A9, blocking its interaction with TLR4 and RAGE receptors. We investigated the efficacy of TASQ in modulating inflammation and improving disease phenotype using *in vitro* and *in vivo* MDS models. Immunofluorescence staining of human BM identified neutrophils and macrophages as primary S100A9 sources. Exposure of mesenchymal stromal cells (MSCs) to S100A9 induced TLR4 downstream signaling, resulting in increased expression of IRAK1, NF-κB-p65, IL-1β, IL-18, caspase 1 and PD-L1. These effects were effectively abolished by TASQ. Additionally, TASQ restored the disturbed MSC-mediated hematopoietic support, as demonstrated by increased numbers of cobblestone area-forming cells and colony-forming units. In NHD13 MDS mice, TASQ (30 mg/kg, 12 weeks) improved hemoglobin and red blood cell counts, but exerted no effect in wild-type (WT) mice. Additionally, TASQ improved bone microarchitecture by increasing trabecular number and bone volume, likely a result of reduced osteoclast activity. Our findings suggest that TASQ mitigates inflammasome activation in the MDS BM, improving erythropoiesis and bone health. These results provide a necessary preclinical basis for clinical trials in lower-risk MDS patients, in whom anemia and osteoporosis often coexist.

## Introduction

Myelodysplastic neoplasms (MDS) are among the most common hematological neoplasms in the elderly population. This group of clonal hematopoietic stem- and progenitor cell (HSPC) disorders are characterized by ineffective hematopoiesis, peripheral cytopenia and a risk for transformation into acute myeloid leukemia (AML). Overall, therapeutic options are limited and allogeneic hematopoietic cell transplantation (allo-HCT) remains the only potential curative treatment option. However, allo-HCT is a burdensome approach associated with numerous complications and is therefore reserved for fit, high-risk patients. Patients with lower-risk (LR) disease, who make up two-thirds of the MDS population, suffer predominantly from anemia and are mostly treated with supportive care (i.e. red blood cell transfusions), erythropoiesis-stimulating agents (ESAs) or luspatercept. These treatments offer only temporary relief and supportive transfusions are associated with impaired quality of life, increased cardiovascular risk, and toxic iron overload^1^. Indeed, disease modification cannot be achieved with these therapies and a relevant proportion of LR-MDS patients will ultimately progress to high-risk MDS or even AML. Therefore, new therapeutic options are urgently needed.

Aberrant innate immune responses and a pro-inflammatory bone marrow (BM) microenvironment play significant roles in MDS pathogenesis. Both, preclinical and clinical data, provide a strong rationale for modulating the disease course by targeting inflammatory pathways, especially in LR-MDS patients^2,3,4^.

The danger-associated molecular pattern (DAMP) molecules S100A8 and S100A9 are key mediators of aberrant inflammation. These molecules bind to Toll-like receptor 4 (TLR4), triggering downstream signaling via IRAK1/4 and TRAF6^3^. This leads to increased NF-κB-mediated transcription of pro-inflammatory cytokines, such as interleukin-1β (IL-1β), tumor necrosis factor α (TNF-α), transforming growth factor β (TGF-β), interleukin-6 (IL-6) and others, which promote further inflammation^5,6^. Additionally, S100A9 can stimulate myeloid-derived suppressor cells (MDSCs) in an autocrine manner via CD33, leading to the secretion of hematopoiesis-suppressive cytokines, such as IL-10 and TGF-β^5^.

Thus, targeting the S100A9 pathway might be an interesting therapeutic strategy to slow-down MDS progression.

Tasquinimod (TASQ, Active Biotech, Lund, Sweden) is a small-molecule oral inhibitor and a second-generation quinolone-3-carboxamide compound that binds to S100A9 and inhibits its interaction with receptors such as RAGE and TLR4^7,8,9^. Originally tested for the treatment of prostate cancer^10^, TASQ has demonstrated immunomodulatory and anti-tumor effects in several preclinical solid cancer models as well as in myelofibrosis^11,12^. The drug is currently being tested in a phase 1 trial in patients with multiple myeloma^13^.

In this study, we investigated whether and how the S100A9-mediated inflammatory activation of the myelodysplastic BM can be rescued by TASQ. We found that mesenchymal stromal cells (MSCs), a central component of the BM microenvironment, are susceptible to adopting an inflammatory phenotype, which causes dysregulation of HSPCs and may contribute to cytopenia in MDS patients. TASQ treatment of MSC cultures *in vitro* mitigated these effects as demonstrated by reduced expression levels of IL-1β, IL-18, caspase 1, and PD-L1 resulting in improved hematopoietic support. Moreover, in NHD13 transgenic mice, TASQ significantly improved hemoglobin levels and red blood cell counts, as well as bone microarchitecture, highlighting its potential as a therapeutic agent for LR-MDS.

## Results

### S100A9 is mainly secreted by neutrophils and macrophages in the human BM

Pro-inflammatory signaling within the BM microenvironment mediated by alarmins, such as S100A9 among others, has been identified as a key pathogenetic driver of MDS^3^. To elucidate the differences in cellular composition and origins of S100A9 within the BM, we undertook a comprehensive approach utilizing multiplex immunofluorescence staining on biopsy sections obtained from patients with LR-MDS and healthy donors. Through the expression of specific markers, including CD271 (MSCs), CD31 (endothelial cells), CD3 (T cells), CD66b (neutrophils), and CD68 (macrophages), we were able to discern the presence and distribution of these cell types in the BM. Intriguingly, S100A9 exhibited high expression in the MDS samples compared to those from healthy individuals (Fig. 1a, b). These findings underscore a distinctive signature within the MDS BM microenvironment.

**Fig. 1.**
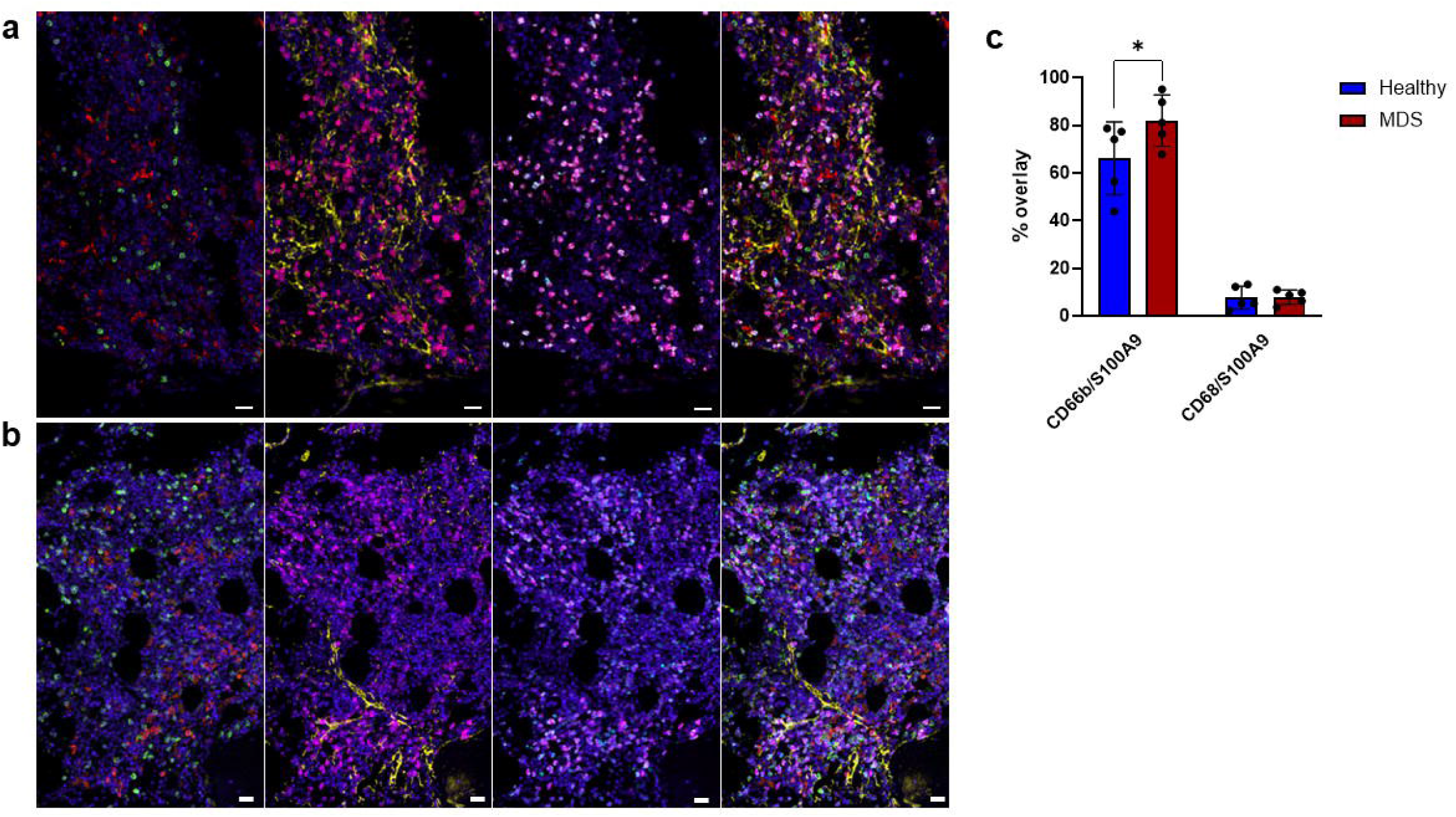
S100A9 is mainly secreted by neutrophils and macrophages in the human BM. Multiplex-immunohistochemistry of BM samples via cyclic immunofluorescence staining by MACSima imaging platform. Representative images of BM tissues from a healthy donor **(a)** and a MDS patient **(b)** are shown, including stainings for CD3^+^ T cells (green) and CD68^+^ macrophages (red); CD271^+^ MSCs (yellow) and S100A9^+^ cells (magenta); CD66b neutrophils (cyan) and S100A9^+^ cells (magenta). In the right images, single stainings are merged. Scale bars indicate 25 µm. **c** The Graph compares the means of S100A9^+^CD66b^+^ neutrophils or S100A9^+^CD68^+^ macrophages between healthy (n=5) donors and MDS patients (n=5), *p< 0.05, by two-way ANOVA with Sidak’s multiple comparisons test.

Co-expression analysis of multiplex immunofluorescence images revealed that the predominant producers and secretors of S100A9 were primarily neutrophils with significant higher intensities in the MDS BM compared to the healthy controls (Fig. 1c). This highlights the pivotal role of these immune cells in the dysregulated expression of S100A9 observed in MDS. Importantly, CD271^+^ MSCs did not express S100A8 or S100A9 or the levels were below the detection limit.

### S100A9 induces a pro-inflammatory signature in MSCs which can be reverted by TASQ *in vitro*

MSCs play a pivotal role within the BM environment, supporting both hematopoiesis and immuno-modulation, as well as serving as precursors for bone-forming cells^14,15^. Thus, we aimed to explore the impact of S100A9 on MSC biology and function. Given the potential for LR-MDS MSCs to lose their inflammatory characteristics *in vitro*, we conducted experiments using recombinant S100A9 to induce inflammatory licensing in both healthy and MDS MSCs, although MDS BM-derived cells might already be “pre-primed”. Since the observed effects were more pronounced in MDS-derived MSCs compared to healthy controls, subsequent data presentation focuses on MDS MSCs unless otherwise specified.

Initially, we investigated S100A9/TLR4-mediated inflammasome activation by examining the expression and activation of downstream molecules through Western blot analysis. Exposure of MSCs to S100A9 induced TLR4 downstream signaling, as evidenced by increased expression of IRAK1 and NF-κB-p65. Notably, the S100A9 inhibitor TASQ reversed the effects and suppressed the expression of these proteins, indicating a reduction in inflammation (Fig. 2a, Suppl. Fig. 1). The expression of TLR4 remained unchanged.

**Fig. 2.**
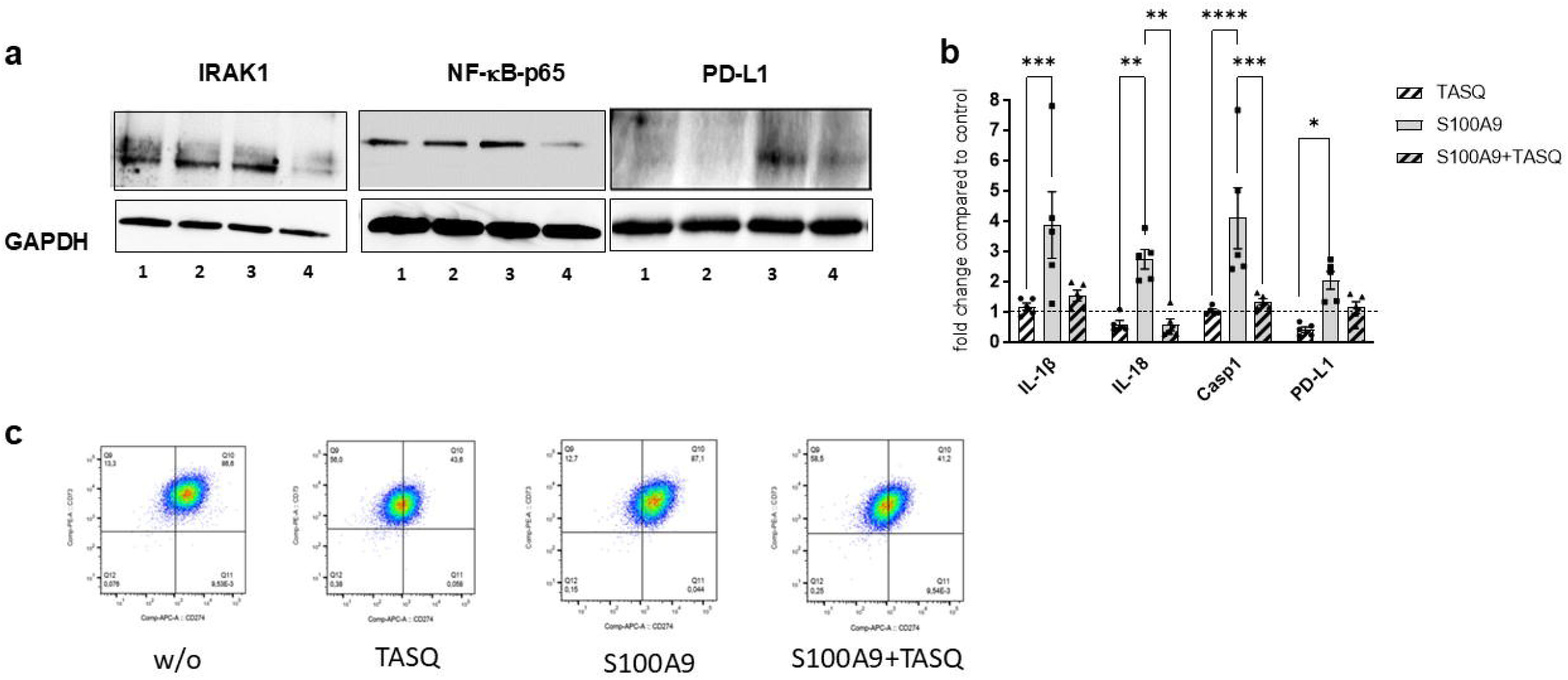
S100A9 induces an inflammatory signature in MDS MSCs which can be reverted by TASQ treatment *in vitro*. MDS and healthy donor MSCs were treated with S100A9 and/or TASQ for three days. **a** Representative Western blot analysis of IRAK1 (80kDa), NF-κB-p65 (65kDa), and PD-L1 (40kDa) performed in MDS MSCs following 72h treatment with recombinant human S100A9 and/or TASQ. GAPDH (37kDa) served as reference protein. The different proteins were analyzed in separate gels and blots. Lane 1: without; lane 2: TASQ; lane 3: S100A9; lane 4: S100A9+TASQ. **b** mRNA quantification of MDS MSCs by real-time PCR. Relative target quantity was determined using the comparative CT (ΔΔCT) method. Amplicons for IL-1β, IL-18, caspase 1 and PD-L1 were normalized to endogenous GAPDH expression and the untreated MSCs were set to 1 (=control). Cumulative data from 5 samples are shown as mean ± SD. Significance was assessed by one-way ANOVA with Tukey’s multiple comparisons test, *p≤ 0.05, **p< 0.01, ***p< 0.001, ****p< 0.0001. **c** Representative flow cytometry plots of MDS MSCs demonstrating increased PD-L1 expression after S100A9 exposure (87.1% positive cells) and reduction after TASQ (43.6% and 41.2% positive cells) treatment.

Additionally, we assessed the mRNA expression levels of potential NF-κB target genes in MSC cultures using quantitative real-time PCR. Our findings revealed a significant increase in mRNA expression of the pro-inflammatory cytokines IL-1β and IL-18, along with their activator caspase 1, in MDS MSCs following treatment with S100A9. Importantly, these effects were attenuated by TASQ (Fig. 2b).

We identified PD-L1 as a potential downstream target of the TLR4/NF-κB signaling pathway in MSCs, where its expression was significantly upregulated by S100A9 stimulation, leading to an approximately 2.5-fold increase compared to baseline levels (Fig. 2b). Notably, this induction of PD-L1 expression could be effectively suppressed by TASQ treatment, reducing it to nearly 50% of the levels observed after S100A9 stimulation. A similar trend was observed at the protein level, where PD-L1 expression was found to be inherently higher in MDS MSCs (Fig. 2a, c, Suppl. Fig 1).

These results demonstrate that S100A9 changes the immune-modulatory function of MSCs and highlight its potential as a therapeutic target as well as the efficacy of TASQ in mitigating the effects.

### MSC differentiation and clonogenic capacity is not significantly modulated by S100A9 and TASQ

The differentiation balance of MSCs is crucial for their hematopoietic support and for balancing bone homeostasis. Therefore, we differentiated cells from MDS patients and healthy donors *in vitro* for 14 days with S100A9/TASQ into the adipogenic and osteogenic lineage. Subsequently, the extent of differentiation was quantified based on the expression of adipogenic- and osteogenic-specific genes, e.g. PPARγ and Runx2. Although the enhanced adipogenic differentiation potential of MDS MSCs could be demonstrated by 36.2-fold vs. 28-fold up-regulation of expression of PPARγ compared to undifferentiated cells, neither S100A9 nor TASQ had a significant influence on these parameters (Suppl. Fig. 2a). Similarly, the osteogenic differentiation capacity of MDS MSCs was not affected by S100A9 or TASQ. Higher mRNA expression of Runx2, a marker for early stages of osteo-chondro-progenitor determination, was detected in MDS compared to healthy MSCs (Suppl. Fig. 2b).

Next, we compared the clonogenic potential of MDS and healthy MSCs, with and without TASQ treatment, by using CFU-F assays. While MDS MSCs formed approximately twice as many colonies as healthy MSCs, treatment with TASQ had no significant effect on MDS MSCs (Suppl. Fig. 2c).

### S100A9 impairs and TASQ restores the supportive function of MSCs towards hematopoiesis *in vitro*

To assess whether S100A9-mediated NF-κB signaling in MDS MSCs contributes to impaired hematopoietic support, we co-cultured CD34^+^ HSPCs from healthy donors on MDS MSC monolayers pre-treated with S100A9 or TASQ. Colony-forming unit assays (CAF-C) were performed weekly for up to four weeks to evaluate hematopoietic support. Notably, S100A9 pre-treatment significantly reduced colony formation, whereas TASQ treatment effectively rescued colony numbers, with significant differences observed at week four (Fig. 3a, b).

**Fig. 3.**
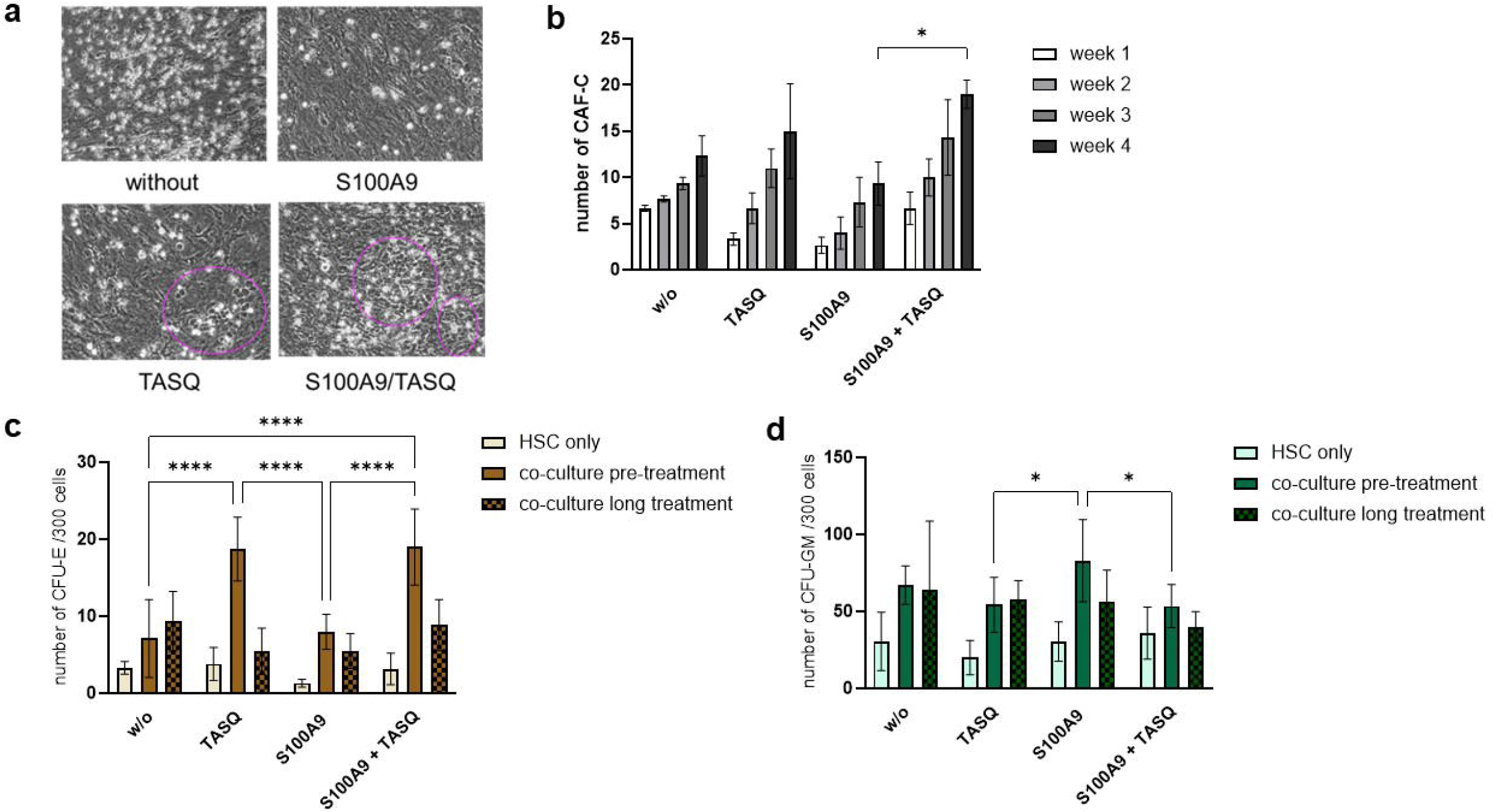
S100A9 impairs and TASQ restores the hematopoietic support function of MSCs *in vitro*. MSCs were pre-treated with S100A9 and/or TASQ for seven days and co-cultured with freshly isolated CD34+ HSPCs. **a** Representative images of CAF-C (highligted with purple circles) for each condition. **b** Quantification of CAF-C after one, two, three and four weeks. Pooled data from N=3 MDS MSC/HSPC co-culture experiments are shown as mean ± SD, *p< 0.05 for S100A9 vs. S100A9+TASQ. **c, d** After one week of co-culture, a CFU assay was performed for 14 days in methylcellulose medium, with or without continuing treatment. Colonies were classified by using the StemVision system. Cumulative data from 4 experiments are shown as mean ± SD, *p< 0.05, ****p< 0.0001 by two-way ANOVA with Tukey’s multiple comparisons test.

To further investigate the impact of MSC treatment on lineage-specific progenitor differentiation, healthy HSPCs were co-cultured on pre-treated MDS MSC layers for one week, followed by an additional two-week colony-forming assay. The colony numbers of erythroid (CFU-E) and granulocyte-macrophage (CFU-GM) progenitors were compared between directly S100A9/TASQ-treated HSPCs and those co-cultured with MSCs pre-treated or continuously exposed to S100A9 or TASQ. HSPCs co-cultured with MSCs exhibited an overall increase in colony formation compared to directly treated cells. S100A9 treatment significantly reduced CFU-E formation in both directly treated HSPCs and those co-cultured with treated MSCs (Fig. 3c), while CFU-GM numbers increased (Fig. 3d). Interestingly, TASQ treatment led to a significant increase in CFU-E numbers exclusively in HSPCs co-cultured with pre-treated MSCs (Fig. 3c), underscoring the supportive role of MSCs during erythroid differentiation. Moreover, TASQ treatment significantly reduced CFU-GM numbers in HSPCs co-cultured with pre-treated MSCs (Fig. 3d), suggesting a potential role in mitigating aberrant myeloid expansion associated with MDS.

### TASQ modulation of MSCs enhances HSC erythroid differentiation

To further investigate the effects of TASQ on MSC-mediated HSC differentiation, healthy BM-derived CD34^+^ cells were co-cultured with MDS MSCs in the presence of erythropoietin (Epo), which is essential for the commitment of erythroid progenitors and their subsequent maturation into mature erythrocytes. To evaluate erythroid differentiation, the expression of CD71 (transferrin receptor), which facilitates iron uptake, and CD235a (glycophorin A), a marker for erythroid precursor and mature erythrocytes, was assessed by flow cytometry.

In the CD34^-^/CD45^-^ fraction, TASQ treatment in the presence of Epo did not alter the percentage of CD71^+^ cells when HSCs were cultured alone. However, a significant increase in CD71^+^ cells was observed in MSC co-culture-derived cells treated with Epo and TASQ (Fig. 4a). In contrast, CD235a expression remained lower, indicating an incomplete maturation of erythroid precursors (not shown). Moreover, the CD34^+^/CD45^+^ fraction was significantly reduced in HSCs derived from TASQ-treated MSC co-cultures compared to untreated co-cultures, to a similar extent as observed with Epo-treated co-cultures. However, neither Epo nor TASQ had a significant effect on this fraction in HSC monocultures (Fig. 4b). These findings highlight the potential role of TASQ in promoting erythroid differentiation through MSC modulation.

**Fig. 4.**
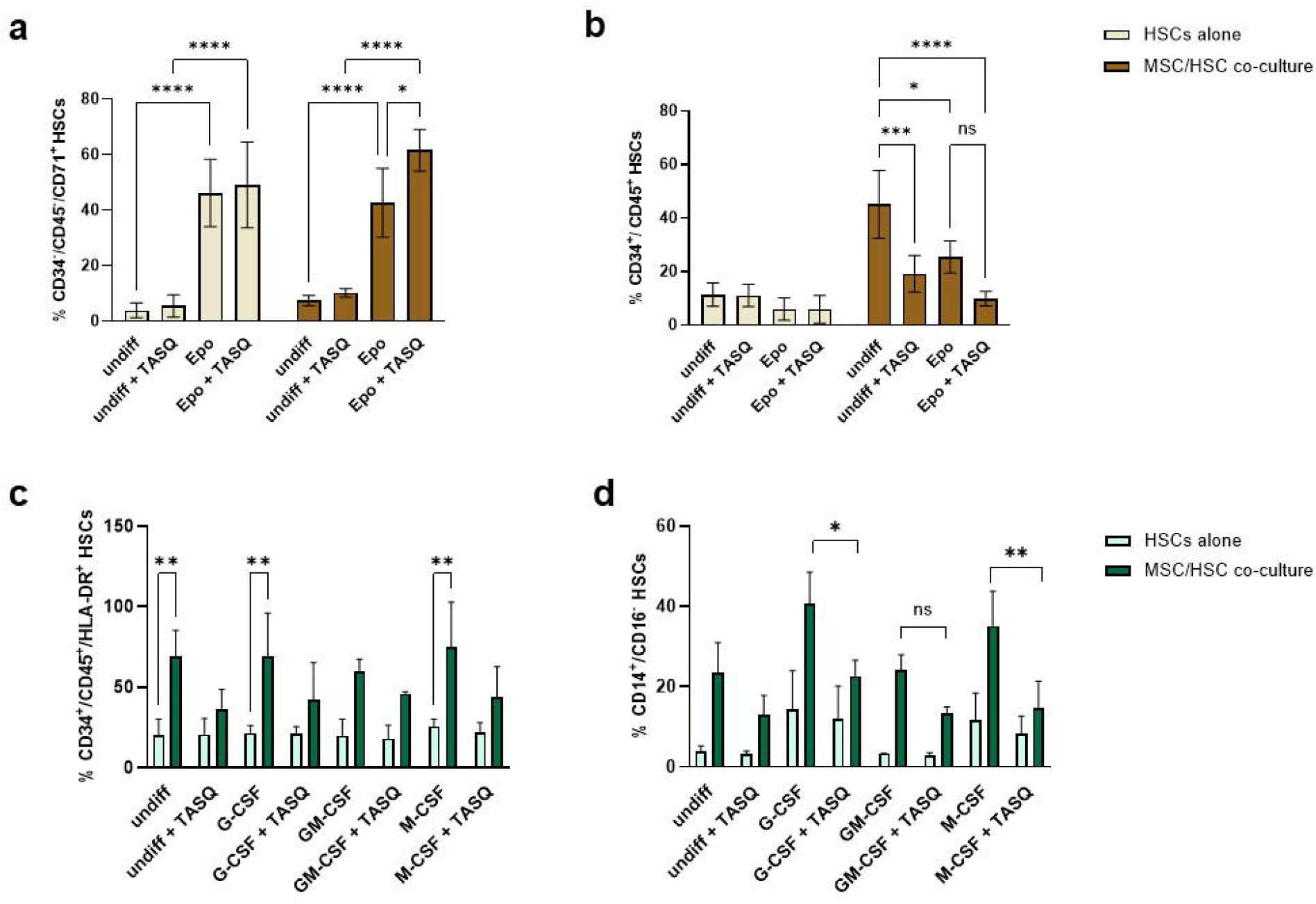
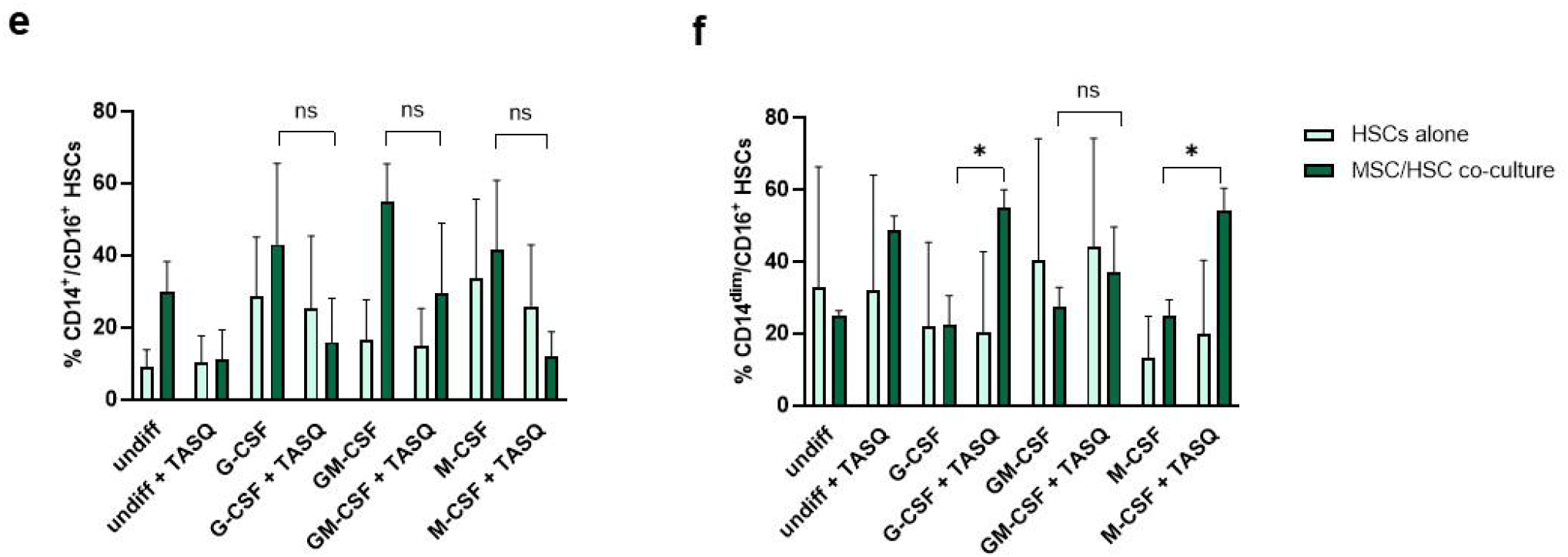
TASQ modulation of MSCs enhances HSC erythroid and myeloid differentiation. MDS and healthy donor MSCs were pre-treated with S100A9 and/or TASQ for seven days and co-cultured with freshly isolated healthy BM CD34^+^ HSCs. **a** Flow cytometry analysis of CD71^+^ erythroid progenitors in the CD34^-^/CD45^-^ fraction shows a significant increase in MSC co-cultures treated with Epo and TASQ. **b** The CD34^+^/CD45^+^ fraction is significantly reduced in TASQ-treated MSC co-cultures, similar to Epo-treated co-cultures, but remains unaffected in HSC mono-cultures. **c** The CD45^+^/CD34^+^/HLA-DR^+^ myeloid progenitor population is significantly elevated in MSC co-cultures, with TASQ reducing its frequency across differentiation conditions. **d-f** Monocyte subset analysis reveals that TASQ decreases CD14^+^/CD16^-^ classical and CD14^+^/CD16^+^ intermediate monocytes in MSC co-cultures, while increasing CD14^dim^/CD16^+^ non-classical monocytes. Data from n=3-4 experiments are shown as mean ± SD, *p< 0.05, **p< 0.01, ***p< 0.001, ****p< 0.0001 by two-way ANOVA with Sidak’s multiple comparisons test.

### TASQ modulation influences myeloid differentiation

To further explore the effects of TASQ on MSC-mediated myeloid differentiation, we assessed specific myeloid progenitor and monocyte subsets under different cytokine conditions.

The proportion of CD45^+^/CD34^+^/HLA-DR^+^ myeloid progenitor cells was significantly increased in co-cultures with MDS MSCs compared to healthy HSCs cultured alone (Fig. 4c). Although not significant, TASQ treatment reduced this population by approximately 25% across all three differentiation conditions, but only in the presence of MSC co-culture (Fig. 4c).

In further analyses of monocyte subset differentiation, CD14^+^/CD16^-^ classical monocytes showed a strong increase in co-culture under all conditions. However, TASQ treatment reduced their numbers by approximately 50% in G-CSF- and M-CSF-driven differentiation, but only in co-cultures (Fig. 4d). Similarly, CD14^+^/CD16^+^ intermediate monocytes exhibited a strong increase in co-culture, with TASQ treatment reducing their numbers by 25–75% across all conditions, again only in co-culture settings (Fig. 4e). Conversely, CD14^dim^/CD16^+^ non-classical monocytes were lower in co-culture compared to mono-cultures, but TASQ treatment increased their proportion by 20–50%, primarily in co-culture conditions (Fig. 4f).

Taken together, our results suggest that TASQ has significant effects on niche-mediated support of erythroid differentiation while mitigating pro-inflammatory myeloid expansion.

### TASQ improves erythropoiesis in NHD13 mice *in vivo*

To investigate whether such effects can be observed *in vivo*, the NHD13 transgenic mouse model was used. The fusion protein in these mice causes a rather slow development of MDS towards high-risk disease and/or AML. At the age of 12 weeks, NHD13 mice already showed mild anemia with decreased red blood cell (RBC) levels (9.5 vs. 10.2×10^12/l, Fig. 5a), prompting treatment initiation at this age. From 12 to 24 weeks, untreated NHD13 mice showed a consistent decline in hemoglobin (Hb) and RBC counts. Both parameters were strongly decreased in 24 weeks old NHD13 mice compared to WT mice (Hb: 7.6 vs. 9.3 mmol/L; RBC: 7.1 vs. 10.2 x10^12^/L). Of note, treatment with TASQ over 12 weeks significantly improved Hb levels to 8.4 mmol/L and RBC counts to 8.2 x10^12^/L in NHD13 mice, while no significant effect was observed in WT mice (Fig. 5a, b).

**Fig. 5.**
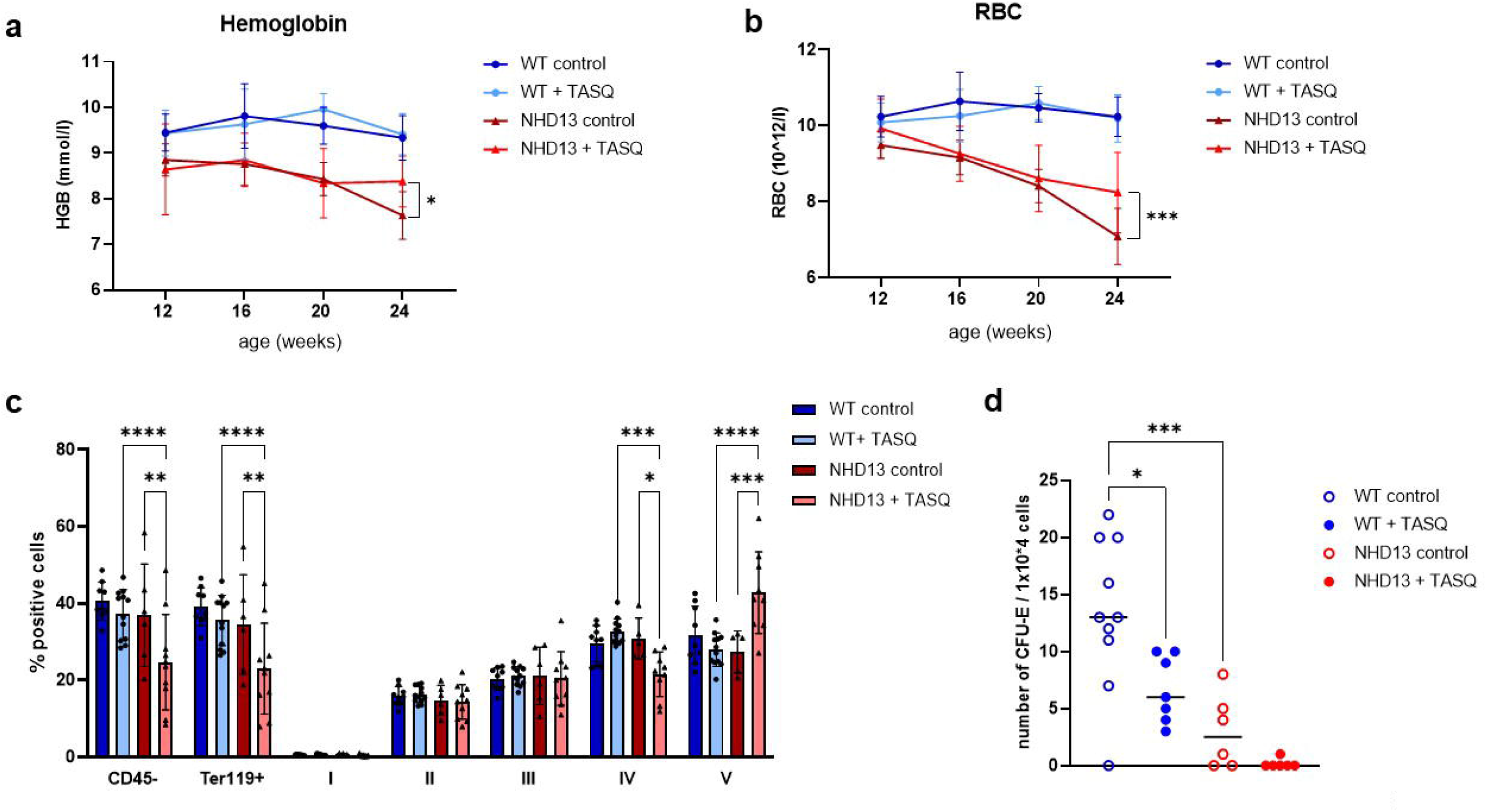
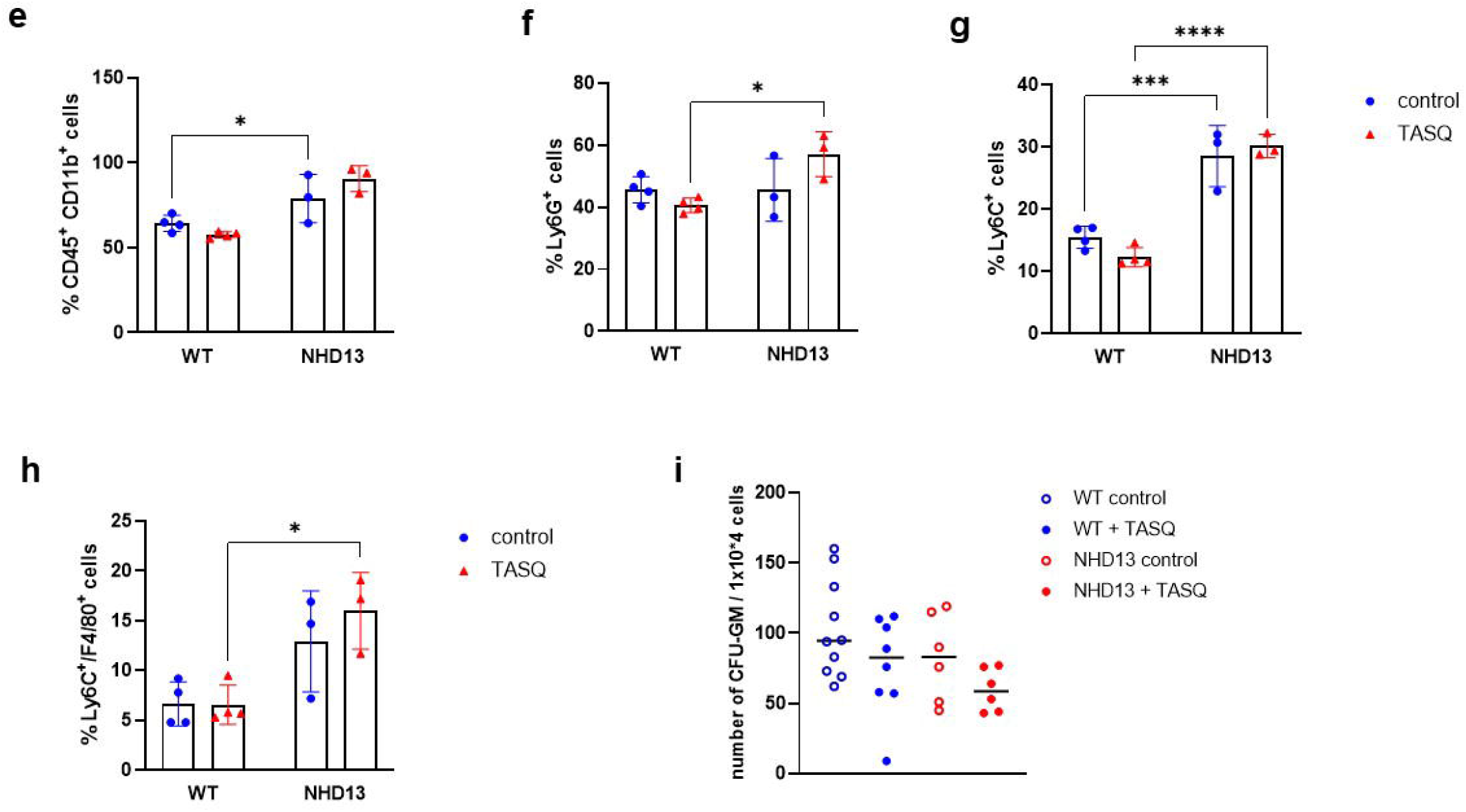
TASQ treatment improves anemia and modulates erythroid and myeloid differentiation in the NHD13 MDS mouse model. Three-month-old NHD13 and WT mice were treated with TASQ-drinking water or normal water over 12 weeks. Peripheral blood cell counts were analyzed every four weeks. **a, b** NHD13 mice exhibited progressive anemia with declining RBC and hemoglobin levels from 12 to 24 weeks. TASQ treatment significantly improved RBC counts and Hb levels in NHD13 mice but had no effect in WT mice. Data from n=10 per group are shown as mean ± SD, *p< 0.05, ***p< 0.001 by two-way ANOVA with Tukey’s multiple comparisons test. **c** Flow cytometry analysis of BM erythroid cells showed a 33% reduction in CD45^−^Ter119^+^ erythroid precursors (stage I-V) in TASQ-treated NHD13 mice, with a shift from late-stage erythroid precursors (stage IV) to reticulocytes (stage V). Data from n=8-10 per group are shown as mean ± SD *p< 0.05, **p< 0.01, ***p< 0.001, ****p< 0.0001 by two-way ANOVA with Tukey’s multiple comparisons test. **d** CFU assays revealed a lower number of erythroid progenitors in NHD13 mice compared to WT controls. TASQ treatment reduced erythroid colony formation in WT mice but had no significant effect in NHD13 mice. Data from n=6-8 are shown as mean ± SD, *p< 0.05, ***p< 0.001 by one-way ANOVA with Tukey’s multiple comparisons test. **e-h** Myeloid BM compartment analysis shows an increased frequency of CD11b^+^ myeloid cells in untreated NHD13 mice, which was not affected by TASQ treatment. Ly6G^+^ granulocytic cells were elevated in TASQ-treated NHD13 mice compared to WT controls, whereas Ly6C^+^ monocytic cells were increased in NHD13 mice but unaffected by TASQ. The Ly6C^+^F4/80^+^ monocyte/macrophage population was expanded in NHD13 mice, with a further slight increase following TASQ treatment. Data from n=3-4 are shown as mean ± SD *p< 0.05, ***p< 0.001, ****p< 0.0001 by two-way ANOVA with Tukey’s multiple comparisons test. **i** CFU assays indicate a reduction in granulocyte-macrophage colony formation in TASQ-treated NHD13 mice, though the decrease was not statistically significant. These findings suggest that TASQ promotes erythroid differentiation while differentially influencing myeloid populations in the BM of NHD13 and WT mice. Data from n=6-8 are shown as mean ± SD.

The platelet count was slightly increased in TASQ treated NHD13 mice at 20 weeks with 673.8 x10^9/L compared to 556.8 x10^9/L in untreated animals, but this difference disappeared at 24 weeks (Suppl. Fig. 3a). The white blood cell (WBC) count was significantly lower in NHD13 than in WT mice but stayed at comparable levels throughout the experiment and was unaffected by TASQ treatment. In contrast, WT mice showed declining WBC levels which were further decreased by TASQ (Suppl. Fig. 3b).

Given the significant improvement observed in peripheral red blood cell counts, we analyzed erythroid precursors in the BM to assess potential changes induced by TASQ treatment (Suppl. Fig. 4a). Notably, the CD45^−^Ter119^+^ population was reduced by 33% in NHD13 mice following TASQ administration (Fig. 5c), suggesting that TASQ promotes erythroid differentiation. Further analysis of the Ter119^+^ compartment revealed an altered distribution of erythroid precursors (stage I-V), characterized by a significant reduction in late-stage erythroid precursors (stage IV) and a concomitant increase in reticulocytes (stage V, Fig. 5c).

To investigate whether these effects were linked to changes in progenitor potential, we performed CFU assays using BM cells. Interestingly, we observed that all colonies of cells from NHD13 were smaller than those of WT mice, with significantly lower numbers of erythroid progenitors. TASQ treatment resulted in a reduction of erythroid colony formation in WT mice, but no significant effect was observed in NHD13 mice (Fig. 5d). These findings suggest that TASQ modulates erythroid differentiation but does not restore erythroid progenitor capacity in the NHD13 model.

Next, we investigated the potential effects of TASQ on the myeloid BM compartment in both NHD13 and WT mice. Flow cytometric analysis (Suppl. Fig. 4b) revealed that the frequency of CD11b^+^ myeloid cells was significantly higher in untreated NHD13 mice compared to WT controls; however, TASQ treatment did not affect their abundance (Fig. 5e). In contrast, Ly6G^+^ granulocytic cells were increased in TASQ-treated NHD13 mice relative to TASQ-treated WT mice (Fig. 5f). Additionally, Ly6C^+^ monocytic cells were significantly elevated in NHD13 mice, and while TASQ treatment led to a reduction in Ly6C^+^ cells in WT mice, it had no effect in NHD13 mice (Fig. 5g). Furthermore, the Ly6C^+^F4/80^+^ monocytic/macrophage population was expanded in NHD13 mice, with a slight but further increase following TASQ treatment, whereas no such effect was observed in WT mice (Fig. 5h). These findings suggest that TASQ differentially influences myeloid cell populations in the BM of NHD13 and WT mice.

Additionally, myeloid progenitors were assessed using CFU assays. While not reaching statistical significance, we observed a decrease in granulocyte-macrophage (GM) colony formation in TASQ-treated NHD13 mice (Fig. 5i), suggesting a potential suppressive effect of TASQ on myeloid progenitor activity.

### Reduced osteoclast number and leads to bone gain in TASQ-treated NHD13 mice

Since NHD13 mice showed impaired bone microarchitecture^16^, we investigated the potential effect of TASQ. Untreated NHD13 mice showed normal bone volume per total volume, but the trabecular number was reduced by 11%, indicating bone loss (Fig. 6b). TASQ treatment for 12 weeks increased the bone volume in both WT and NHD13 cohorts (WT and NHD13: 2-fold), accompanied by an increase in trabecular number (WT: +29%; NHD13: +32%; Fig. 6a, b). To evaluate the cause of changes in the bone remodeling, bone histomorphology of the femora and vertebrae was performed and the bone turnover markers were assessed in the serum. Surprisingly, the number of bone-forming osteoblasts and their activity was not increased by TASQ, and the latter was even reduced by 37% in WT mice (Fig. 6c, d). However, TASQ treatment led to a reduction in the bone formation rate (WT: -42%; NHD13: -51%) and osteoid surface per bone surface (WT: -66%; NHD13: -75%; Fig. 6e, f). Furthermore, a reduced number of osteoclasts (−42%) was detected in NHD13 mice (Fig. 6g). TASQ treatment led to a reduction in the number of osteoclasts of about 53% in WT mice only. However, the osteoclast activity was significantly decreased in both WT and NHD13 mice as measured by TRAcP5b levels in the serum (WT: -20%; NHD13: -26%; Fig. 6g, h). Thus, the reduced bone resorption predominantly led to an increase in bone volume. Furthermore, a higher maximal force was detected in TASQ-treated WT mice in the compression test of vertebrae (+60%), while NHD13 mice had already an elevated value (+35%), which was not affected by TASQ (Fig. 6i).

**Fig. 6.**
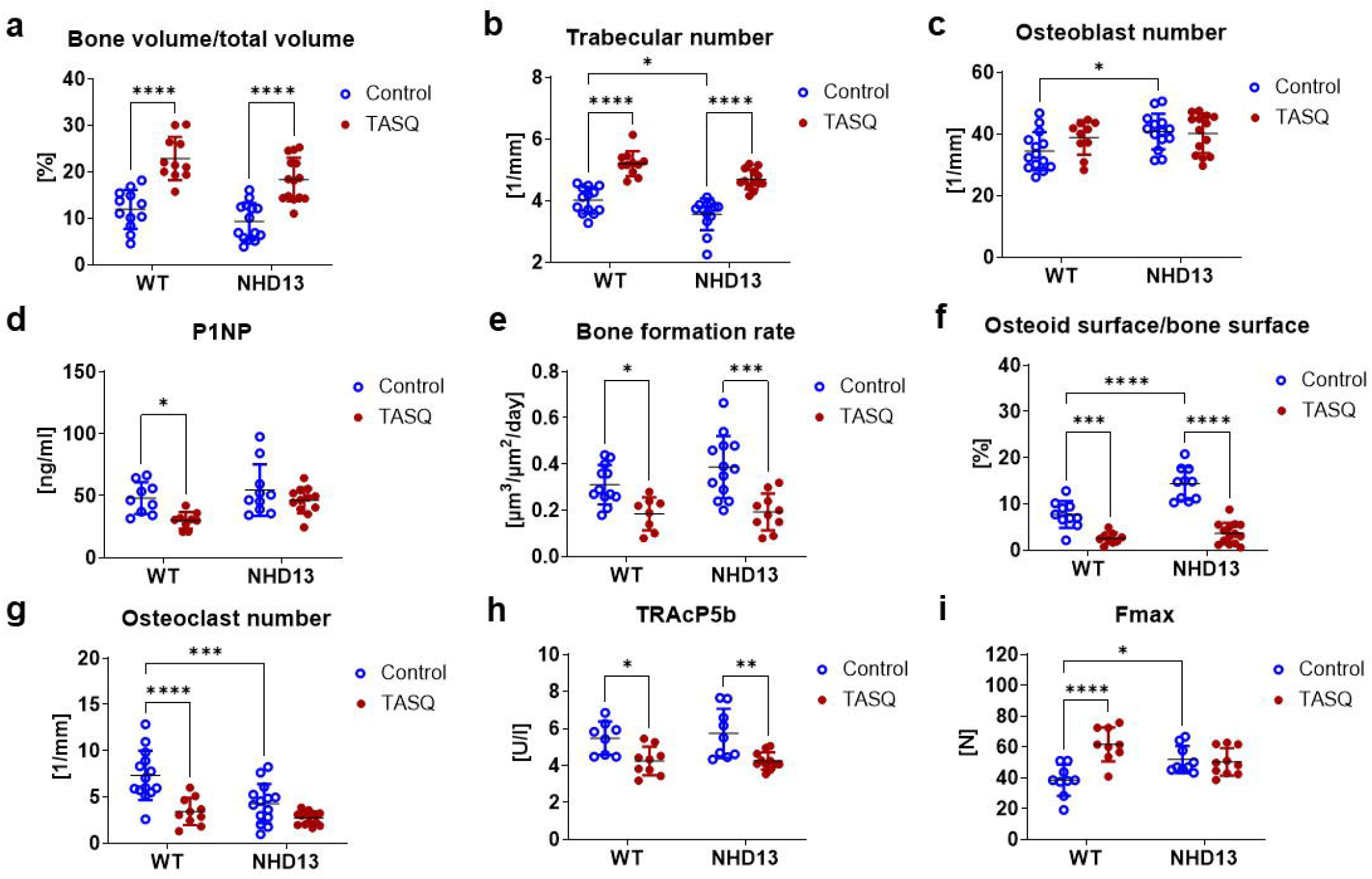
TASQ treatment improves bone microarchitecture of NHD13 mice. Three-month-old NHD13 and WT mice were treated with TASQ-drinking water or normal water over 12 weeks. After treatment femora/vertebrae were used to analyze the bone microarchitecture and phenotype. Using micro computed tomography **(a)** the bone volume (n=11-14) and **(b)** the trabecular number (n=11-13) in the femora were assessed. Afterwards, femora were stained with Tartrate-resistant acid phosphatase (TRAP) and vertebrae were used for von Kossa/van Gieson staining and for the analysis of the calcein double labeling. TRAP-staining were used to determine **(c)** the number for osteoblasts (n=11-14) and **(d)** their activity was measured in the serum with P1NP ELISA (n=9-12). **e** Based on the calcein labeling, the bone formation rate was assessed in vertebral bodies (n=8-13). **f** The osteoid surface per bone surface were analyzed in von Kossa/van Gieson stained vertebrae (n=9-14). **g** TRAP-stained femora were used to determine osteoclast number (n=10-14) and (H) their activity was assessed by TRAPc5b serum levels (n=8-11). **i** The fifth lumbar vertebrae were used for the compression test to measure the maximal force (n=9-10). Data are shown as mean ± SD. Statistical analysis was performed by two-way ANOVA for the effect of MDS, TASQ treatment, and the interaction of both. Statistical significance of Bonferroni multiple comparisons is denoted. *p<0.05; **p<0.01; ***p<0.001; ****p<0.0001.

Taken together, TASQ treatment enhances bone volume and strength in both WT and NHD13 mice by reducing bone turnover.

## Discussion

MDS is characterized by cytopenia and an increased risk of progression to AML. These features are driven in part by dysregulation of the BM microenvironment, shifting from a pro-inflammatory state in early disease stages to an immunosuppressive milieu in later stages^4,16,17,18^. Current treatments are limited, highlighting the need for new therapies. Fromowitz et al. identified TLR4 and TLR6 as highly overexpressed in MDS, with TLR4 inhibition shown to promote erythroid and myeloid maturation *in vitro*^19^. The TLR4 ligands S100A8/A9 correlate with anemia severity^20^, suggesting a potential therapeutic target. We therefore conducted a comprehensive preclinical study to assess the effects of the S100A9-inhibiting small molecule TASQ on the MDS BM, both *in vitro* and *in vivo*.

We and others identified neutrophils and, to less extent, macrophages as the predominant source of S100A9 in the human BM^20,21^. While CD271^+^ MSCs do not appear to be directly involved in S100A9 production, they undergo significant modulation through the influence of extracellular S100A9, such as cellular senescence^22^. This process involves the induction of the TLR4 receptor downstream signaling, formation of the NLRP3 inflammasome, and subsequent secretion of IL-1β. These findings suggest that elevated levels of S100A9 in the BM microenvironment could contribute to stromal cell aging and dysfunction^22^. Transcriptome analysis of CD271^+^ niche cells in a prospective, homogeneously treated cohort of LR-MDS patients revealed that expression of S100A8 and S100A9 was strongly correlated with a subgroup of MDS patients demonstrating significant overexpression of this heterodimer independent of established prognostic factors and activation of p53 and TLR programs in mesenchymal cells, in line with experimental data from a mouse model pointing at the existence of a p53-S100A8/9-TLR axis^23^. The controversial data on the relationship between S100A9 and MSCs underline the enormous heterogeneity of MSC subpopulations and complex regulatory mechanisms.

We demonstrated that exposure of both MDS and healthy MSCs to S100A9 triggered a TLR4-dependent downstream signaling cascade *in vitro*, resulting in increased expression of IRAK1, NF-kB-p65, Caspase 1, and the pro-inflammatory cytokines IL-1β and IL-18. TASQ treatment effectively dampened the S100A9-induced inflammatory response of MSCs.

One mechanism to suppress inflammation and dampen the immune response is governed by the immune checkpoint receptor PD-1 and its ligand PD-L1. Cheng et al. reported that S100A9 contributes to ineffective hematopoiesis in MDS via induction of PD-1 and PD-L1 expression^24^. In this context, the increased death of CD34^+^ HSPCs and CD71^+^ erythroid progenitor cells play a key role. In line with these data, we detected a significant higher expression of PD-L1 mRNA and protein in MSCs after S100A9 exposure which was effectively suppressed by TASQ treatment. Moreover, effective downregulation of PD-L1 in MDS-derived MSCs has been associated with enhanced hematopoietic support^24^, a finding that was also reflected in our *in vitro* experiments. Our results with HSPC/MSC co-cultures align with previous reports indicating that S100A9 plays a critical role in MDS pathophysiology by promoting chronic inflammation and disrupting the BM microenvironment^5,24^. The observed reduction in CAF-C numbers following S100A9 treatment supports prior evidence that TLR4/NF-κB pathway activation in MSCs negatively impacts their hematopoietic support function^25,26^. This impairment may contribute to the ineffective hematopoiesis observed in MDS patients.

TASQ has been recognized for its immune-modulating properties, particularly in cancer and inflammatory conditions^11^. Our results indicate that TASQ rescues hematopoietic support by MSCs, potentially by counteracting NF-κB-mediated dysfunction with potential involvement of PD-L1 as an identified target in these conditions. These findings are consistent with previous studies demonstrating that modulating NF-κB signaling in MSCs can restore their supportive role in hematopoiesis^14^. The lineage-specific effects observed in our study further emphasize the impact of S100A9 on hematopoietic differentiation. The reduction in CFU-E formation following S100A9 treatment is in line with previous reports that inflammatory signals suppress erythropoiesis^27^. Conversely, the increased CFU-GM numbers suggest a shift towards myeloid expansion, a hallmark of MDS^28^. The ability of TASQ to restore CFU-E numbers while reducing CFU-GM highlights its potential role in rebalancing hematopoiesis.

Several studies have demonstrated that MSCs provide critical signals for lineage commitment and differentiation^14,28^. Our findings suggest that TASQ modulates MSCs in a way that promotes erythroid differentiation while reducing primitive HSC fractions, indicating a shift toward lineage commitment. The observed increase in CD71^+^ cells in TASQ-treated MSC co-cultures suggests an enhanced early erythroid commitment. This is consistent with prior reports indicating that additional erythropoietic factors and microenvironmental cues are required for complete erythroid maturation^30^.

The observed reduction in CD34^+^/CD45^+^ HSCs in TASQ-treated co-cultures suggests a shift away from self-renewal and toward lineage commitment. Similar findings have been reported in studies where MSCs influence HSC fate through niche signaling^31^. This supports the notion that MSCs, when properly modulated, can act as key regulators of hematopoietic lineage specification. Importantly, the decrease in CD34^+^ progenitor frequency in our study aligns with reports showing that microenvironmental modulation can promote efficient erythroid differentiation while reducing the pool of undifferentiated progenitors^32^.

In the NHD13 murine model, TASQ had a significant impact on red blood cell counts and clonogenic capacity, confirming our previous *in vitro* results. However, in the murine system, the regulatory mechanisms underlying these results may vary depending on the specific mouse model used. A detailed BM analysis revealed lower erythroid progenitor colonies in treated mice which is in contrast to the CFU assays performed with isolated human MSCs and HSPCs demonstrating increased erythroid colony number after TASQ treatment. A potential alternative regulatory mechanism is that TASQ enhances erythroid maturation and survival, thereby increasing erythrocyte output from a smaller progenitor pool. Additionally, TASQ may suppress ineffective erythropoiesis by enhancing the differentiation efficiency of erythroid precursors or reducing apoptosis at later stages of maturation^33^. Although TASQ does not increase erythropoiesis to WT levels in NHD13 mice, it is evident that more severe anemia is prevented. A shift in lineage dynamics, such as accelerated progression through the CFU-E stage, could also explain the reduced colony numbers despite increased RBC production.

The expansion of CD11b^+^ myeloid cells and Ly6C^+^ monocytic populations in untreated NHD13 mice is consistent with the myeloid bias and ineffective differentiation characteristic of MDS^34^. TASQ treatment did not reverse these abnormalities but selectively modulated specific subsets, such as increasing Ly6G^+^ granulocytic cells and slightly expanding the Ly6C^+^F4/80^+^ population. The reduction in GM colony formation following TASQ treatment in NHD13 mice suggests a potential suppressive effect on myeloid progenitor activity, which may be linked to its differentiation-promoting effects on the erythroid lineage. Prior studies have highlighted the complex interplay between erythroid and myeloid differentiation, with skewed lineage commitment being a hallmark of MDS^35^. The differential effects of TASQ in WT versus NHD13 mice suggest it modulates progenitor activity, but its impact is context-dependent. Further studies are warranted to elucidate the mechanistic basis of these effects.

TASQ treatment not only modulated hematopoiesis by the BM microenvironment, but also improved bone microarchitecture, including increased trabecular number and bone mass, and reduced osteoclast activity in NHD13 mice. This result is supported by our human *in vitro* co-culture of MSCs and HSCs differentiated to monocytes, the precursors of osteoclasts. Since pro-inflammatory cytokines such as IL-6, IL-1β, and TNF-α enhance bone resorption, the inhibitory effects of TASQ likely contribute to bone preservation and the mitigation of bone loss in MDS patients^36^. An increase in trabecular bone volume with TASQ has also been described in multiple myeloma but the authors did not mentioned whether the osteoclasts are responsible for the effect^37^.

While our findings provide novel insights into the effects of TASQ on hematopoiesis and bone microarchitecture, the study has potential limitations. First, although the inhibition of the S100A9/TLR4/NF-κB axis appears to play a relevant role, the precise mechanistic pathways through which TASQ influences MSC-mediated hematopoiesis remain unclear. Second, the long-term functional consequences of TASQ treatment on immune homeostasis and erythropoiesis remain to be determined. While we observed significant changes in erythroid and myeloid differentiation, it is unclear whether these effects persist over time and may even have the potential to delay disease progression. Longitudinal studies are warranted to assess whether prolonged TASQ treatment alters immune function, clonal evolution, or overall hematopoietic stability. Finally, although NHD13 mice exhibit key features of LR-MDS and are a widely accepted murine model for the disease, they only partly reflective of human MDS pathophysiology. In particular, differences in inflammasome activation, disease progression, and inflammatory mediators may limit the direct translational applicability of our findings. Additionally, the timing and duration of TASQ treatment could significantly impact its effects, necessitating further investigations to optimize treatment strategies and better align with human MDS pathophysiology. In summary, our data provide compelling evidence that TASQ can attenuate pathological S100A9/TLR4/NF-κB-mediated inflammasome activation in the myelodysplastic BM niche resulting in improved hematopoietic support by MSCs *in vitro*. These findings highlight the potential of TASQ as a therapeutic agent for MDS patients, who often suffer from concomitant anemia and bone loss. Of note, the therapeutic effects of TASQ in MDS are not solely attributed to remodeling of the BM niche, as cell-intrinsic effects have also been described by other groups. Notably, Schneider et al. identified significant expression of S100A8/9 in mutant erythroblasts, leading to a p53-mediated defect in erythroid differentiation, which could be reversed by genetic inactivation of S100A8^21^. However, our study is the first to elucidate the niche-specific effects of TASQ in MDS and their broader impact on the disease phenotype. Given the limited success of various anti-inflammatory therapies in the past (e.g., Canakinumab trial, IRAK4 inhibitors), we propose that a S100A9-targeted anti-inflammatory approach, which intervenes early in the inflammatory cascade, presents a promising therapeutic strategy. Further investigation, particularly in clinical trials, is warranted to fully evaluate the therapeutic potential of TASQ in MDS.

## Materials and Methods

### Patients

MSCs were collected from hematologically healthy donors undergoing hip replacement (n= 4, age 63-82 years), BM stem cell donors (n=5, age 23-44), and untreated patients with MDS (n=12, age 41-80 years, patient’s characteristics are depicted in Suppl. Tab. 1) under written informed consent (according to the Declaration of Helsinki) as part of the MDS registry (EK289112008) and the BoHemE Study (NCT02867085) at the University Hospital in Dresden. Heparinized BM samples were obtained at diagnostic aspiration or during hip total arthroplasty surgery.

### Human BM immunofluorescence staining

To analyze the bone marrow using immunofluorescence, 3–5 μm thick paraffin sections were used. Deparaffinization, rehydration, and epitope retrieval were done manually via an established protocol. Briefly, sections were deparaffinized in xylene (2 × 15 min, VWR International, Fontenay-sous-Bois, France) and hydrated by washes of graded ethanol (Berkel AHK, Ludwigshafen, Germany) to water (B. Braun, Melsungen, Germany). Tissue sections were boiled in TRIS-HCL buffer (Akoya biosciences, Akoya Biosciences, Menlo Park, California, USA) at pH 9.0 for 20 min for antigen retrieval. Multiplex-immunohistochemistry of bone marrow sections was performed via cyclic immunofluorescence staining on MACSima imaging platform (Miltenyi Biotec, Bergisch-Gladbach, Germany). The procedure is based on cyclic immunofluorescence staining. Every sequence includes: staining the tissue with fluorophore-coupled antibodies (Suppl. Tab. 2), microscope the region of interest (ROI), and eliminating the fluorescence signal by bleaching. The staining is performed with commercially available antibodies that are directly linked to a fluorophore. Three dyes and DAPI are measured in one cycle with minimal spectral overlap. The staining cycles are conducted in the fully automated system of MACSima Imaging Platform. Analysis software MACSiQView was used for image data evaluation, just for nuclei segmentation was performed with a StarDist 2D plugin in ImageJ^38^.

### Cell culture

BM mononuclear cells were collected using a Ficoll gradient and cultured at 37°C and 5% CO_2_ in Dulbecco’s modified Eagle’s medium (DMEM, Invitrogen) supplemented with 10% FCS. After 24 h, non-adherent cells were removed, and adherent cells were expanded. Primary MSC from passages 2–5 were used for the experiments. Sub-confluent cultures were treated with 1.5 µg/ml S100A9 (Abcam) and 10 µM TASQ (provided by Active Biotech) for up to one week. Cell growth and proliferation were calculated by cell counting with trypan blue.

HSPCs were isolated using CD34 antibody-conjugated magnetic beads, according to the manufacturer’s instructions (Miltenyi Biotec). The purity of the isolated CD34+ population was confirmed by flow cytometry and only fractions >95% positivity were used in the analysis.

### Clonogenic assays

Cobblestone area forming-cell (CAF-C) assays were performed over 4 weeks using pre-treated healthy or MDS MSC layers. One thousand magnetically isolated CD34^+^ cells were added in LTC-IC medium (Stem Cell Technologies) supplemented with 1×10^−6^ M hydrocortisone (Sigma Aldrich). Colony forming unit (CFU) assays were carried out using cells harvested after one week of coculture, with 1×10^4^ cells being plated in enriched methylcellulose medium with recombinant cytokines (MethoCult H4435, Stem Cell Technologies). Colonies were counted after 2 weeks and classified under a microscope or with the StemVision system (Stem Cell Technologies).

### Mice

For the *in vivo* experiments, 3-month-old heterozygous NHD13 (C57BL/6-Tg(Vav1-NUP98/HOXD13)G2Apla/J) and their littermate wild-type (WT) mice were used. The NUP98-HOXD13 (NHD13) fusion gene has been detected in patients with MDS and mice expressing the NHD13 fusion gene under the vav1 promotor develop MDS-like symptoms and BM alterations^39,40^. All mouse experiments were approved by the institutional animal care committee as well as the Federal State of Saxony (TVV 64/2022). Mice were fed a standard diet as well as water ad libitum, and they were exposed to a 12-hour light/dark cycle in an air-conditioned room at 23°C. To analyze the effects of TASQ on hematopoiesis and bone homeostasis, NHD13 and WT mice were treated with normal water (control) or TASQ water (30 mg/kg per body weight) for 12 weeks. The blood count was monitored at the baseline and every 4 weeks until the end of the experiment using the XN-1000 (Sysmex, Norderstedt, Germany). All mice received an intraperitoneal calcein injection (20 mg/kg body weight) two and five days before sacrifice. Finally, long bones, spine, bone marrow, and serum were collected for subsequent analysis. One femur and the spine per mouse were fixed in 4% PFA for 48 h and stored in 50% ethanol.

The NUP98 gene encodes a protein that is involved in RNA and protein transport across the nuclear membrane^40^. Fusion genes with NUP98 have been identified in various hematologic malignancies, including MDS.

### Flow cytometry

Flow cytometry was applied to phenotypically characterize HSPCs after isolation and co-culture experiments. The used antibodies are depicted in table S4. Furthermore, the BM of NHD13 and WT mice was flushed with PBS supplemented with 5% fetal calf serum (FCS; Biochrom) as well as 5 mM EDTA, and cells were stained with an antibody cocktail consisting of anti-murine antibodies (Suppl. Tab. 4). Data were evaluated using FlowJo 10.10.0 software.

### Mouse BM serum analysis

Concentrations of the bone turnover markers procollagen type 1 amino-terminal propeptide (P1NP) and tartrate-resistant acid phosphatase 5b (TRAcP5b) were measured in serum using ELISA according to the manufacturers’ protocols (IDS, Frankfurt/Main, Germany).

### Bone microarchitecture

For analysis of the bone microarchitecture, μCT (vivaCT40, Scanco Medical) was performed on excised femora with an isotropic voxel size of 10.5 μm (70 kVP; 114 μA; 200 ms integration time). We used established protocols from Scanco Medical to evaluate the trabecular bone parameters and analyzed 100 slices beneath the growth plate.

### Bone histomorphometry and bone strength

The lumbar vertebrae (L1-5) were isolated for further analysis. L1/L2 were embedded in paraffin and 2 µm sections were used for TRAP staining to determine the number of osteoclasts, osteoblasts, and osteocytes per bone perimeter. Von Kossa/van Gieson staining was performed on 4 µm methyl methacrylate section of L3/L4 to evaluate bone mineralization. In addition, 7 µm sections of L3/L4 were used to assess the distance between the two calcein labeling lines by fluorescence microscopy. All analyses were performed with the Osteomeasure software and pictures were taken with the CellSens program or the AxioVision 4.8 program.

To determine the vertebral stiffness, first, L5 was transferred to PBS one day before compression testing (Zwick Roell, Ulm, Germany). The L5 was placed onto the lower plate and the mechanical force was applied vertically onto the vertebra via the upper plate. The maximal load (Fmax) was measured and the bone stiffness was calculated using testXpert II—V3.7 software (Zwick Roell, Ulm, Germany).

## Statistical analysis

Statistical analysis was performed using GraphPad Prism software version 9.5.0 (GraphPad Software). Data are presented as mean ± standard deviation (SD). All experiments were repeated at least three times. For multiple group comparisons, two-way ANOVA with post hoc Tukey’s, Sidak’s, or Bonferroni test was performed. A p-value of less than 0.05 was regarded as being statistically significant.

## Additional methods

Additional methods are described under supplemental information.

## Acknowledgments

The authors would like to thank Katrin Müller, Ivonne Habermann, Claudia Richter, Claudia Dill, and Robert Kuhnert for excellent technical support.

## Author contributions

Conceptualization: MW, HW, MT, EV, HE, EB, MB, KS

Methodology: MW, HW, RW, KM, ALB, TD, KS

Investigation: HW, ALB, KM, TD, RW

Visualization: MW, ALB, KM, HW, RW

Funding acquisition: MW, KS, MB

Project administration: MW, KS

Interpretation: MW, HW, RW, ALB, KM, TC, MB, KS

Supervision: MB, UP, MR, LCH

Writing – original draft: MW, HW, KS, RW

Writing – review & editing: MW, HW, RW, ALB, KM, EB, MT, EV, HE, SW, UP, TC, LCH, MR, MB, KS

## Data and materials availability

All data are available in the main text or the supplementary materials.

---

**Fig. 7.**
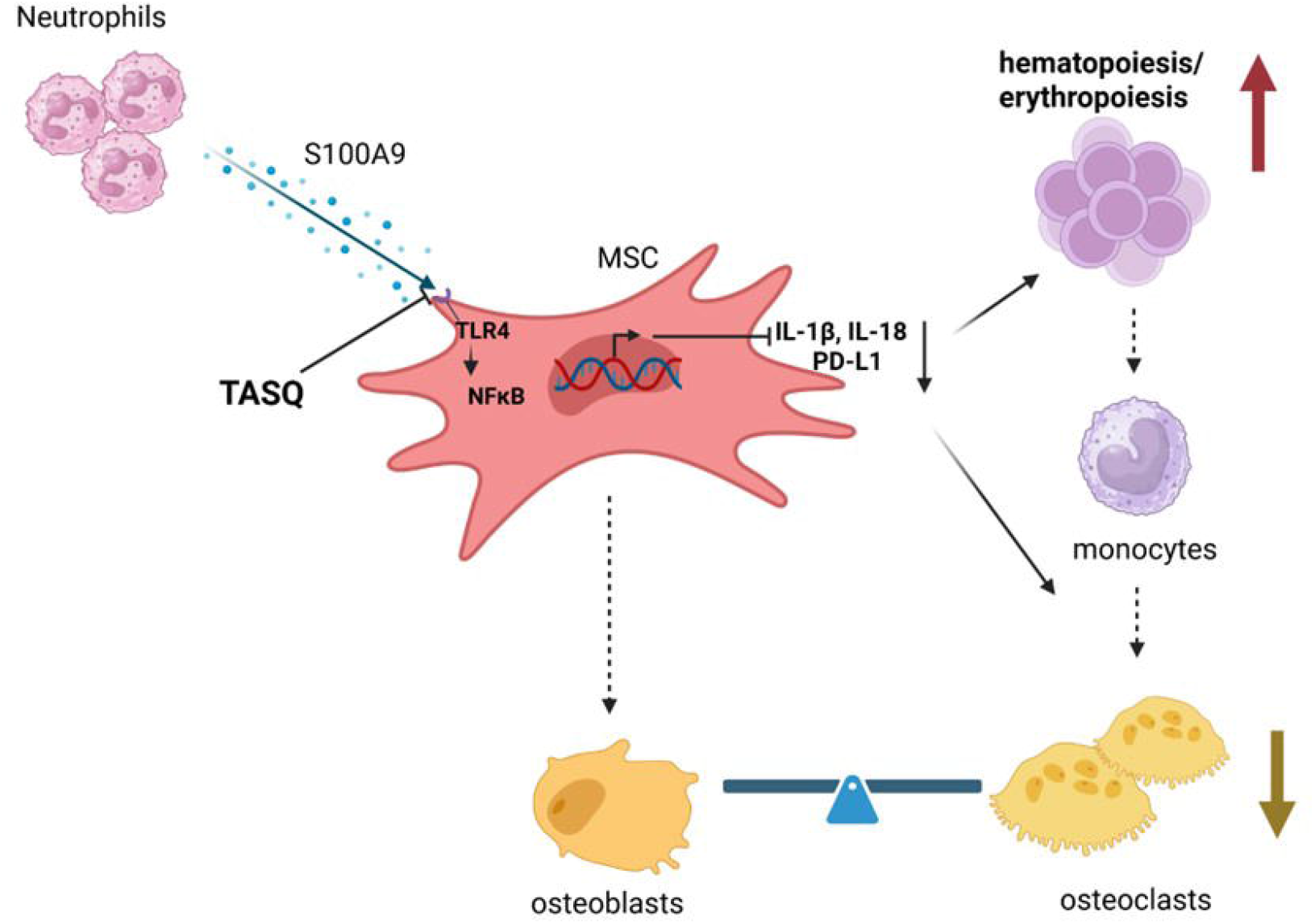
Graphical summary. (generated with bioRender). Neutrophil-derived S100A9 activates TLR4/NF-κB signaling in MDS MSCs, inducing pro-inflammatory cytokines (e.g., IL-1β, IL-18) and PD-L1 expression. TASQ treatment attenuates this inflammatory response, leading to reduced cytokine and PD-L1 levels. This promotes improved hematopoietic support and erythropoiesis, modulates monocyte maturation, reduces osteoclast activity, and restores bone homeostasis.

