## Supplemental material for "Preclinical Efficacy of Tasquinimod in Myelodysplastic Neoplasms: Restoring Erythropoiesis and Mitigating Bone Loss"

**Supplementary Materials**

**Materials and Methods**

**Western blot**

Whole cell lysates were prepared using RIPA lysis buffer containing proteinase inhibitors. Protein concentrations were determined using the BCA Protein Assay Kit (Thermo). Equal protein amounts were separated by SDS-polyacrylamide gel electrophoresis and transferred to polyvinylidene ﬂuoride membranes (Bio-Rad Laboratories) for detection of speciﬁc proteins using primary antibodies for IRAK1, NFkBp65, PD-L1 and GAPDH (all from Cell Signaling) overnight at 4°C, following secondary antibodies HRP-conjugated goat anti-mouse IgG (Invitrogen) or donkey anti-rabbit IgG (GE Healthcare), respectively (Suppl. Tab. 3). The blots were incubated with ECL Plus Western Blotting reagent (Amersham) and the signals captured using a LAS3000 imaging system. Band intensity was quantified by ImageJ software. All Western blot experiments were run in triplicates to ensure reproducibility and reliability of the data.

**Real-time polymerase chain reaction (RT-PCR)**

RNA was isolated from MSC	using Trizol (Thermo) and reverse transcribed into cDNA using RevertAid cDNA synthesis kit (Thermo) with oligo-dT primers. Relative target quantity was determined using the comparative CT (∆∆CT) method. RT-PCR was performed using SYBR Green/ROX PCR master mix (Thermo) and target specific primers (Tab. S5) on a Quantstudio 3 cycler (Applied Biosystems). Amplicons were normalized to endogenous GAPDH control.

**MSC differentiation assays**

MSCs in passage 2 were seeded in 6-well plates (5 × 10^3^ MSCs/cm^2^), cultured to subconfluency for approximately 4 days in DMEM, and subjected to adipogenic (0.5 mM 1-methyl-3-butylisoxanthine, 1 μM dexamethasone, 100 μM indomethacin, 10 μM insulin) or osteogenic (0.1 μM dexamethasone, 0.2 mM ascorbate-2-phosphate, 10 mM b-glycerophosphate) differentiation in the presence or absence of S100A9/TASQ with medium exchange twice a week. After 14 days, the cells were collected for mRNA isolation, and quantitative real-time PCR was performed for adipo- (PPARγ) and osteo-specific (Runx2) genes.

**Colony forming unit-ﬁbroblast (CFU-F) assay**

MSCs in passage 2 were seeded in 6-well plates (2x10^2^/ well) with 3 ml of MSC Expansion Media (Miltenyi Biotec). TASQ was added to MDS MSCs. No medium exchange was performed. At day 14, supernatant was removed. The cell layer was washed twice with water, ﬁxed with methanol and stained with Giemsa’s azur eozin methylen blue-solution (Merck) for 30 min. Afterwards, the colonies were counted. Each assay was performed in triplicate.

**Figure captions**

**Fig. S1. Densitometric quantification of Western blot bands.** Western blot analysis of IRAK-1, NF-kB-p65 and PD-L1 was performed for MSCs from MDS patients after the respective treatment and the band intensity was quantified by Image J software. Cumulative data from 3 samples are shown as mean ± SD. Significance was assessed by two-way ANOVA with Tukey’s multiple comparisons test, *p≤ 0.05, **p< 0.01, ***p< 0.001, ****p< 0.0001.

**Fig. S2. MSC differentiation and clonogenic capacity is not significantly modulated by S100A9 and TASQ.** MDS and healthy donor MSCs were treated with S100A9 and/or TASQ during the respective experiments. (A) MSCs were incubated for 14 days in adipogenic differentiation media, the mRNA was isolated and the expression of PPARγ was quantified by real-time PCR. (B) MSCs were incubated for 14 days in osteogenic differentiation media, the mRNA was isolated and the expression of Runx2 was quantified by real-time PCR. Relative target quantity was determined using the comparative CT (∆∆CT) method and amplicons were normalized to endogenous GAPDH expression and the undifferentiated MSC samples were set to 1 (=control). (C) For Colony-forming unit fibroblast (CFU-F) assays, cells were cultured in MSC Expansion medium for 14 days without exchange, MDS MSCs in the presence or absence of TASQ, and colonies were counted after Giemsa staining.

**Fig. S3. TASQ treatment had no significant influence on platelets and white blood cells in NHD13 MDS mice.** Three-month-old NHD13 and WT mice were treated with TASQ-drinking water or normal water over 12 weeks. Peripheral blood cell counts were analyzed every four weeks. (A) NHD13 mice exhibited lower platelet levels than WT animals. TASQ treatment led to a slight, non-significant increase from week 16 onwards. The white blood cell (WBC) count was significantly lower in NHD13 than in WT mice but stayed at comparable levels throughout the experiment and was unaffected by TASQ treatment. WT mice showed declining WBC levels which were significantly decreased by TASQ in week 16 and 20. Data from n=10 per group are shown as mean ± SD, ***p< 0.001, ****p< 0.0001by two-way ANOVA with Tukey’s multiple comparisons test.

**Fig. S4. Gating strategy for murine BM analysis.** (A) Erythroid precursors were assessed by staining with antibodies against CD45, Ter119, and CD44. (B) Myeloid precursors were assessed with antibodies against CD45, CD11b, Ly6C, Gr1, and F4/80.

**Table S1. Patient data**

| **No.** | **Diagnosis WHO 2022** | **Sex** | **Age** | **Karyotype** | **Molecular genetics (mutation; VAF%)** | **IPSS** | **IPSS-R** |
| --- | --- | --- | --- | --- | --- | --- | --- |
| 1 | MDS with SF3B1 | m | 69 | 46,XY[14] | SF3B1 mutation | NA | low |
| 2 | MDS with SF3B1 | f | 79 | 46,XX[3] | DNMT3A (p.P307R; 46.3%), MPL (p.Y591C; 8.6%), SF3B1 (p.K700E; 46.9%) | low | int |
| 3 | MDS-LB MLD | f | 75 | 46,XX[7] | ASXL1 (p.Leu823fs; 40.61%), SRSF2 (p.Pro95His; 37.57%), TET2 (p.Gly1169Arg; 37.88%), TET2 (c.3500+1G>A; 40.86%), TET2 (p.Met533fs; 1.44%) | low | low |
| 4 | MDS-LB MLD | m | 58 | 46,XY[16] | U2AF1 (p.Ser34Tyr; 21%) | int-1 | low |
| 5 | MDS-LB MLD | m | 54 | NA | ASXL1 (p.Glu635fs; 23.55%), CEBPA (p.Ala274fs; 8.24%), CEBPA (p.Lys276fs; 8.89%), RUNX1 (p.Arg204*; 9.75%), RUNX1 (p.Arg320*; 4.31%), RUNX1 (p.Ala475fs; 1.49%), SRSF2 (p.Pro95His; 47.17%), STAG2 (p.Arg861fs; 26.71%), TET2 (p.Tyr1560fs; 33.05%), TET2 (p.Arg1216*; 45.69%) | NA | low |
| 6 | MDS-LB SLD | m | 75 | 46,XY,del(20)(q11.2)[17]/46,XY[4] | RUNX1 (p.Arg201Gln; 50%), SRSF2 (p.Pro95Arg; 47%), TET2 (p.Met695fs; 51%), TET2 (p.Gln654*; 47%), TP53 (p.Arg282Trp; 37%) | low | int |
| 7 | MDS-LB MLD | f | 70 | 46,XX[17] | no detectable mutations | int-1 | low |
| 8 | MDS/MPN with SF3B1 mutation and thrombocytosis | f | 74 | NA | SF3B1 (p.Lys700Glu; 41.51%), MPL (p.Trp515Ser; 4.62%), TP53 (p.Ser94Pro; 2.17%) | int-1 | int |
| 9 | MDS-IB1 | m | 71 | 46,XY[20] | DDX41 (p.Lys331del; 48.38%), DDX41 (p.Asp344Glu; 14.94%), DNMT3A (p.Leu737fs; 0.5%), GATA2 (p.Asp46Asn; 49.15%) | NA | int |
| 10 | MDS-IB1 | m | 76 | 46,XY[24] | CUX1 (p.Pro1249Argfs*71; 36%), STAG2 (p.Glu511Ter; 95%), TET2 (p.Pro851Leufs*22; 11%), SRSF2 (p.Pro95_Arg102del; 63%) | int-1 | int |
| 11 | MDS-IB1 | m | 43 | 46,XY[24] | no detectable mutations | low | high |
| 12 | MDS-IB1 | m | 59 | 47,XY,+21[19] | ASXL1 (p.Glu635fs; 9.24%), CBL (p.Arg420Gln; 10.4%), NF1 mutations (0.08%, 1.9%, 4.8%) | high | very high |
| 13 | no known hematologic disease | f | 44 | NA | NA |  |  |
| 14 | no known hematologic disease | m | 32 | NA | NA |  |  |
| 15 | no known hematologic disease | m | 23 | NA | NA |  |  |
| 16 | no known hematologic disease | m | 42 | NA | NA |  |  |
| 17 | no known hematologic disease | m | 43 | NA | NA |  |  |
| 18 | no known hematologic disease | f | 63 | NA | NA |  |  |
| 19 | no known hematologic disease | f | 82 | NA | NA |  |  |
| 20 | no known hematologic disease | f | 68 | NA | Non-CHIP |  |  |
| 21 | no known hematologic disease | m | 68 | NA | NA |  |  |

**Table S2. Antibodies for immunofluorescence**

| **Antigen** | **Clone** | **Company** |
| --- | --- | --- |
| **CD3** | REA1151 | Miltenyi biotec, Bergisch Gladbach, Germany |
| **CD68** | REA1305 | Miltenyi biotec, Bergisch Gladbach, Germany |
| **CD271** | REAL709 | Miltenyi biotec, Bergisch Gladbach, Germany |
| **CD66b** | REA306 | Miltenyi biotec, Bergisch Gladbach, Germany |
| **CD34** | QBEnd10 | Abcam, Cambridge, UK |
| **S100A9** | D5O6O | Cell Signaling, Danvers Massachusetts, USA |

**Table S3. Antibodies for Western blot**

| **Antigen** | **Clone** | **Host/Isotype** | **Dilution** | **MW (kDa)** | **Source** |
| --- | --- | --- | --- | --- | --- |
| **NF-kB- p65** | L8F6 | Mouse IgG2b | 1:1000 | 65 | Cell Signaling, Danvers Massachusetts, USA |
| **IRAK1** | D51G7 | Rabbit IgG | 1:1000 | 78-105 | Cell Signaling, Danvers Massachusetts, USA |
| **PD-L1** | E1L3N | Rabbit IgG | 1:1000 | 40-50 | Cell Signaling, Danvers Massachusetts, USA |
| **GAPDH** | 14C10 | Rabbit IgG | 1:1000 | 37 | Cell Signaling, Danvers Massachusetts, USA |

**Table S4. Flow cytometry antibodies and settings for measurement at FACSymphony A3**

|  |  |  |  |  |  |  |  | **Settings** | | |
| --- | --- | --- | --- | --- | --- | --- | --- | --- | --- | --- |
| **Antigen** | **Fluoro-chrome** | **Clone** | **Host** | **Reactivity** | **Subtype** | **Source** | **Dilution** | **Laser** | **Filter** | **PMT** |
| **human HSC** |  |  |  |  |  |  |  |  |  |  |
| HLA-DR | FITC | L243 | mouse | human | IgG2a,k | BD Biosciences | 25 | 488F | 530/30 | 320 |
| CD45 | V500 (PO) | HI30 | mouse | Human |  | BD Biosciences | 20 | 405G | 525/50 | 350 |
| CD14 | PE | M5E2 | mouse | human | IgG2a,k | BD Biosciences | 25 | 561 D | 586/15 | 490 |
| CD34 | APC | AC136 | mouse | human | IgG2a,k | Miltenyi Biotec | 40 | 637C | 670/30 | 407 |
| CD34 | PE | AC136 | mouse | Human | IgG2a,kappa | Miltenyi | 20 | 561 D | 586/15 | 490 |
| CD16 | APC-H7 | 3G8 | mouse | human | IgG1,k | BD Biosciences | 20 | 637A | 780/60 | 450 |
| CD235a | FITC | 10F7MN | mouse | human | IgG1, kappa | eBiosciences | 20 | 488F | 530/30 | 320 |
| CD71 | APC | L01.1 | mouse | human | IgG2a | BD Pharmingen | 25 | 637C | 670/30 | 407 |
| DAPI |  |  |  |  |  |  |  | 355G | 450/50 | 180 |
| **mouse BM cells** |  |  |  |  |  |  |  |  |  |  |
| GR1 (Ly-6C/G) | FITC | RB6-8C5 | rat | mouse | IgG2b,k | BioLegend | 200 | 488 F | 530/30 | 400 |
| LY6C | PE | HK1.4 | rat | mouse | IgG2c,k | eBioscience | 160 | 561 D | 586/15 | 380 |
| CD45 | PE-Cy7 | 30-F11 | rat | mouse/human | IgG2b,k | eBioscience | 200 | 561 A | 780/60 | 420 |
| F4/80 | APC | BM8 | rat | mouse | IgG2a,k | eBioscience | 20 | 637C | 670/30 | 500 |
| CD11b | A700 | M1/70 | rat | mouse | IgG2b,k | eBioscience | 80 | 637B | 730/45 | 470 |
| CD34 | FITC | RAM34 | rat | mouse/rat/human | IgG2b,k | eBioscience | 80 | 488 F | 530/30 | 400 |
| CD38 | PE | 90 | rat | mouse | IgG2c,k | eBioscience | 160 | 561 D | 586/15 | 450 |
| Ter119 (Ly-76) | FITC | TER-119 | rat | mouse/human | IgG2b,k | eBioscience | 100 | 488 F | 530/30 | 317 |
| CD44 | PE | IM7 | rat | mouse/human | IgG2a,k | eBioscience | 200 | 561 D | 586/15 | 380 |
| CD3 | FITC | 17A2 | rat | mouse | IgG2b,k | BD Bioscience | 200 | 488 F | 530/30 | 350 |
| CD19 | FITC | eBio1D3 | rat | mouse/ human | IgG2b,k | eBioscience | 1000 | 488 F | 530/30 |  |
| NK1.1 | FITC | PK136 | rat | mouse/ human | IgG2c,k | eBioscience | 300 | 488 F | 530/30 |  |
| Ter119 (Ly-76) | FITC | TER-119 | rat | mouse/human | IgG2b,k | eBioscience | 400 | 488 F | 530/30 |  |
| CD11b | FITC | M1/70 | rat | mouse | IgG2a,k | BioLegend | 1000 | 488 F | 530/30 |  |
| GR1 (Ly-6C/G) | FITC | RB6-8C5 | rat | mouse | IgG2b,k | BioLegend | 1000 | 488 F | 530/30 |  |
| CD45R (B220) | FITC | RA3-6B2 | rat | mouse/ human | IgG2b,k | eBioscience | 150 | 488 F | 530/30 |  |
| Sca-1 (Ly-6A/E) | PE-Cy5 | D7 | rat | mouse/ human | IgG2a,k | eBioscience | 200 | 561 B | 670/30 | 420 |
| CD117 (c-kit) | APC-A780 | 2B8 | rat | mouse/human | IgG2b,k | eBioscience | 200 | 637 A | 780/60 | 500 |
|  | DAPI |  |  |  |  |  |  | 355G | 450/50 | 250 |
|  | PI |  |  |  |  |  | 100 | 561 C | 610/20 | 350 |

**Table S5. Primer used for real-time PCR**

| human | IL-18 | Forward | ACTGCCTGGACAGTCAGCAA |
| --- | --- | --- | --- |
|  |  | Revers | GCAGCCATCTTTATTCCTGAGA |
| human | IL-1β | Forward | CTCTTCGAGGCACAAGGCAC |
|  |  | Revers | CAAGTCATCCTCATTGCCACTGT |
| human | CASP1 | Forward | TGAGCAGCCAGATGGTAGAGC |
|  |  | Revers | TCACTTCCTGCCCACAGACAT |
| human | PD-L1 | Forward | GGCATCCAAGATACAAACTCAA |
|  |  | Revers | CAGAAGTTCCAATGCTGGATTA |
| human | PPAR-γ | Forward | CGAGAAGGAGAAGCTGTTGG |
|  |  | Revers | TCAGCGGGAAGGACTTTATG |
| human | Runx2 | Forward | GCCTTCAAGGTGGTAGCCC |
|  |  | Revers | CGTTACCCGCCATGACAGTA |
