## Supplementary figures and images for "Preclinical Efficacy of Tasquinimod in Myelodysplastic Neoplasms: Restoring Erythropoiesis and Mitigating Bone Loss"

### Suppl. Fig. 1

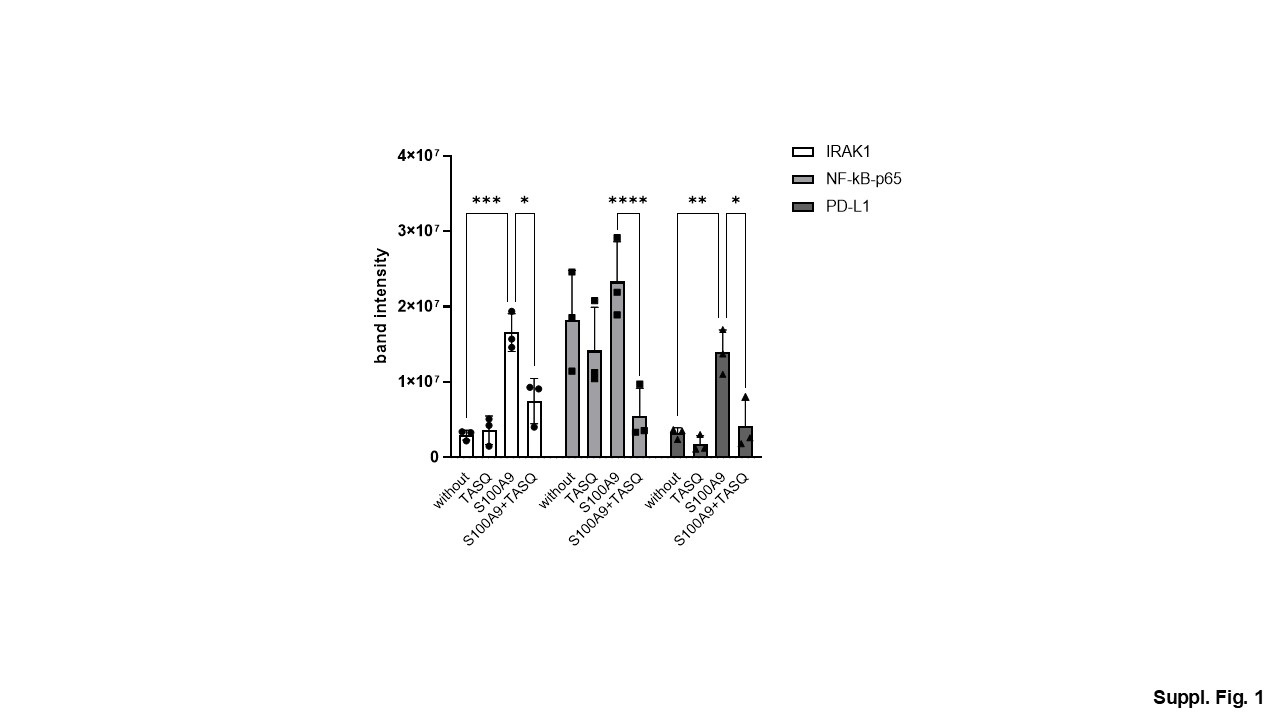

### Suppl. Fig. 2

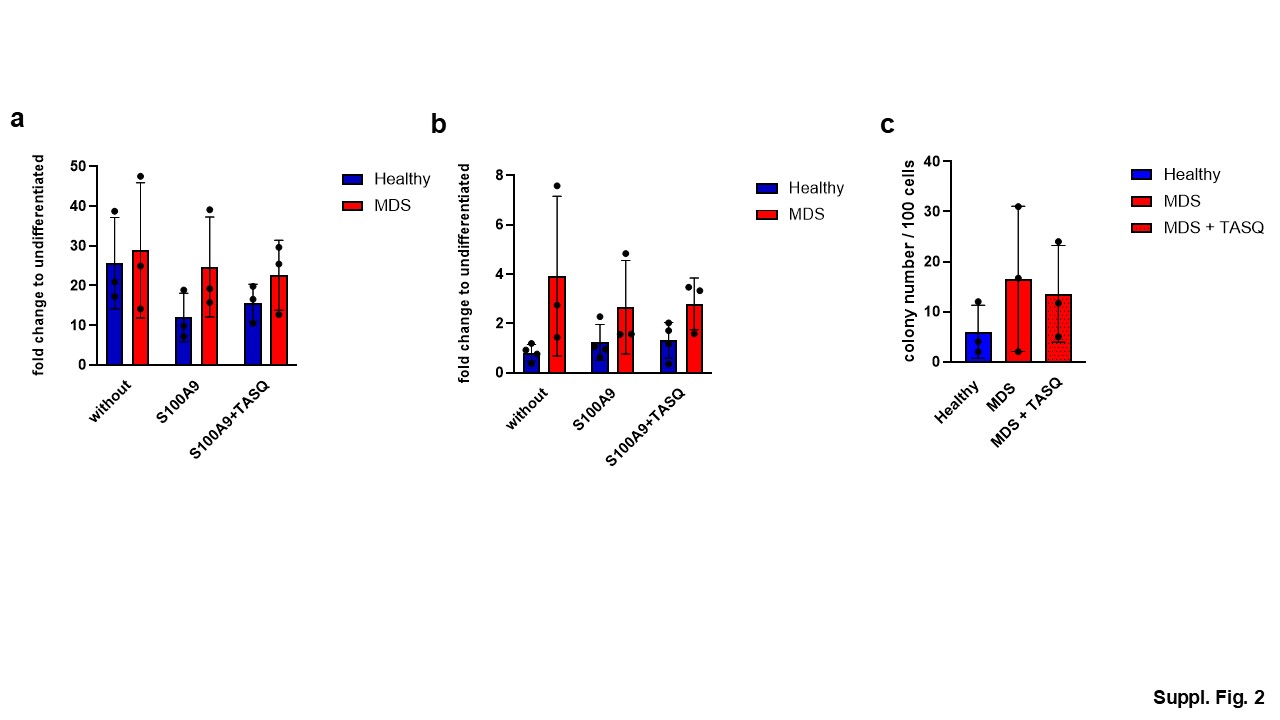

### Suppl. Fig. 3

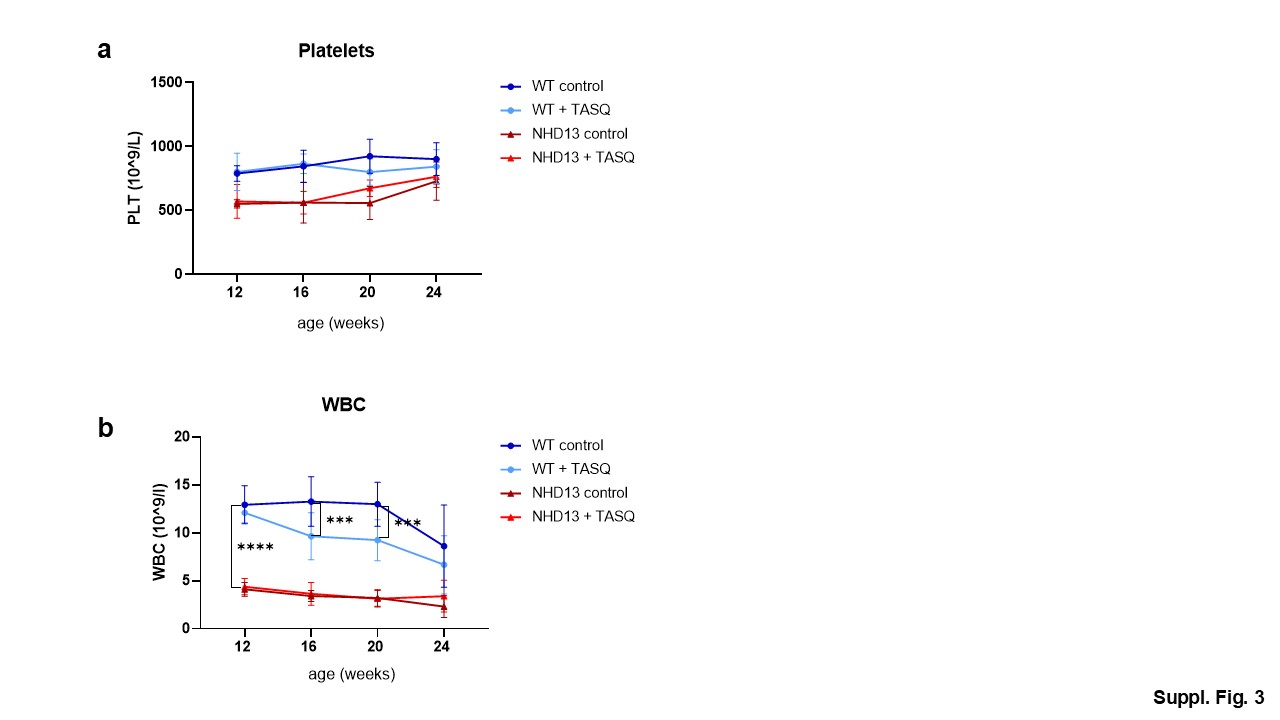

### Suppl. Fig. 4a

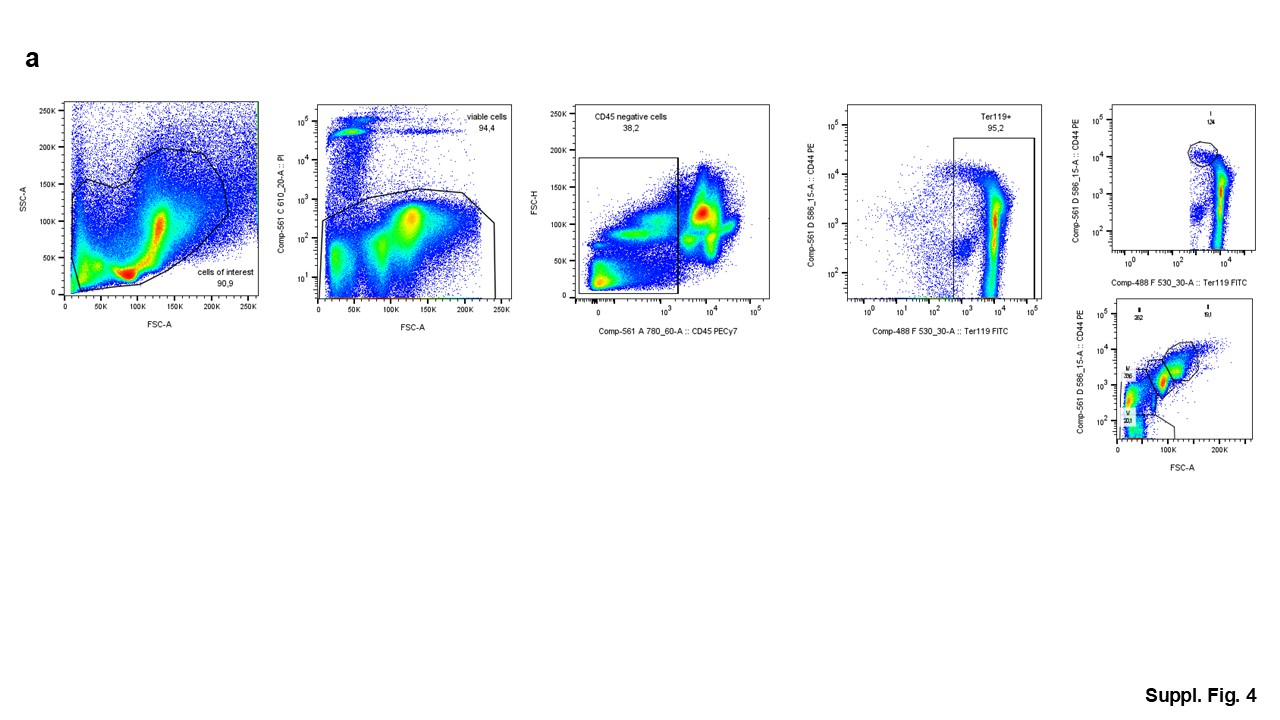

### Suppl. Fig. 4b

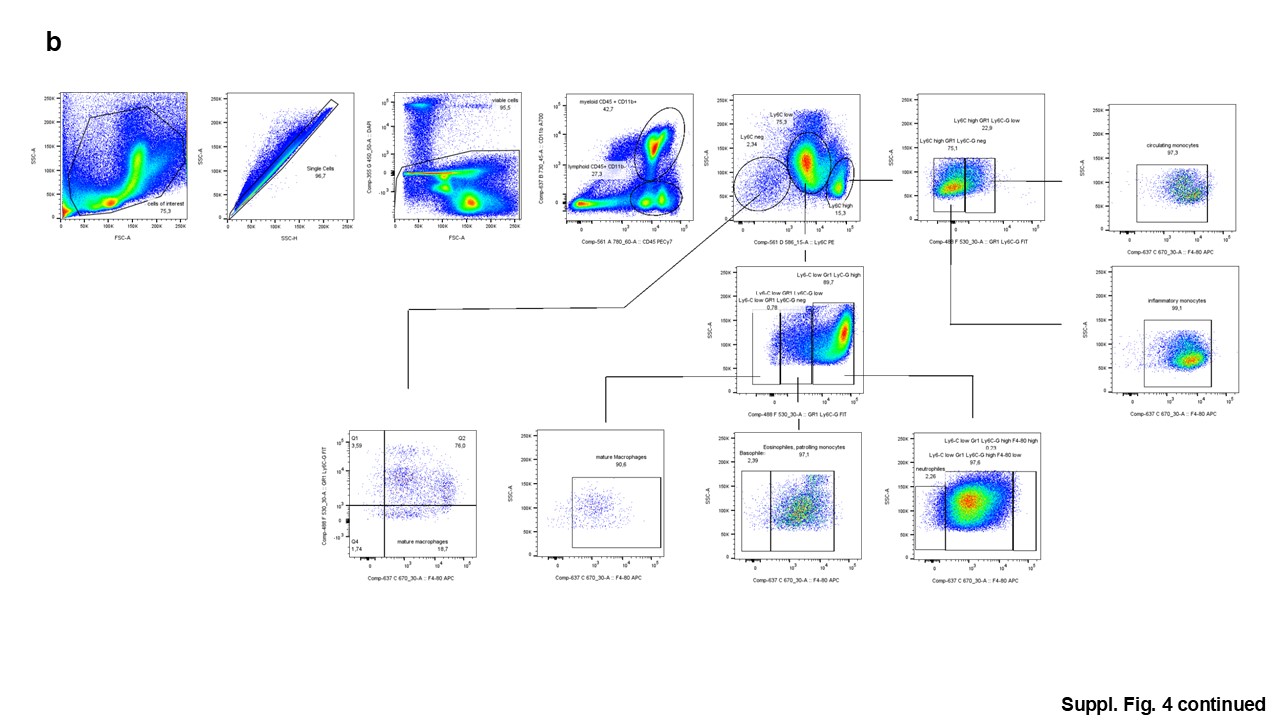
